## Supplementary_Figures for "Single cell spatial transcriptomics integration deciphers the morphological heterogeneity of atherosclerotic carotid arteries": Supplementary_figures_1.pdf

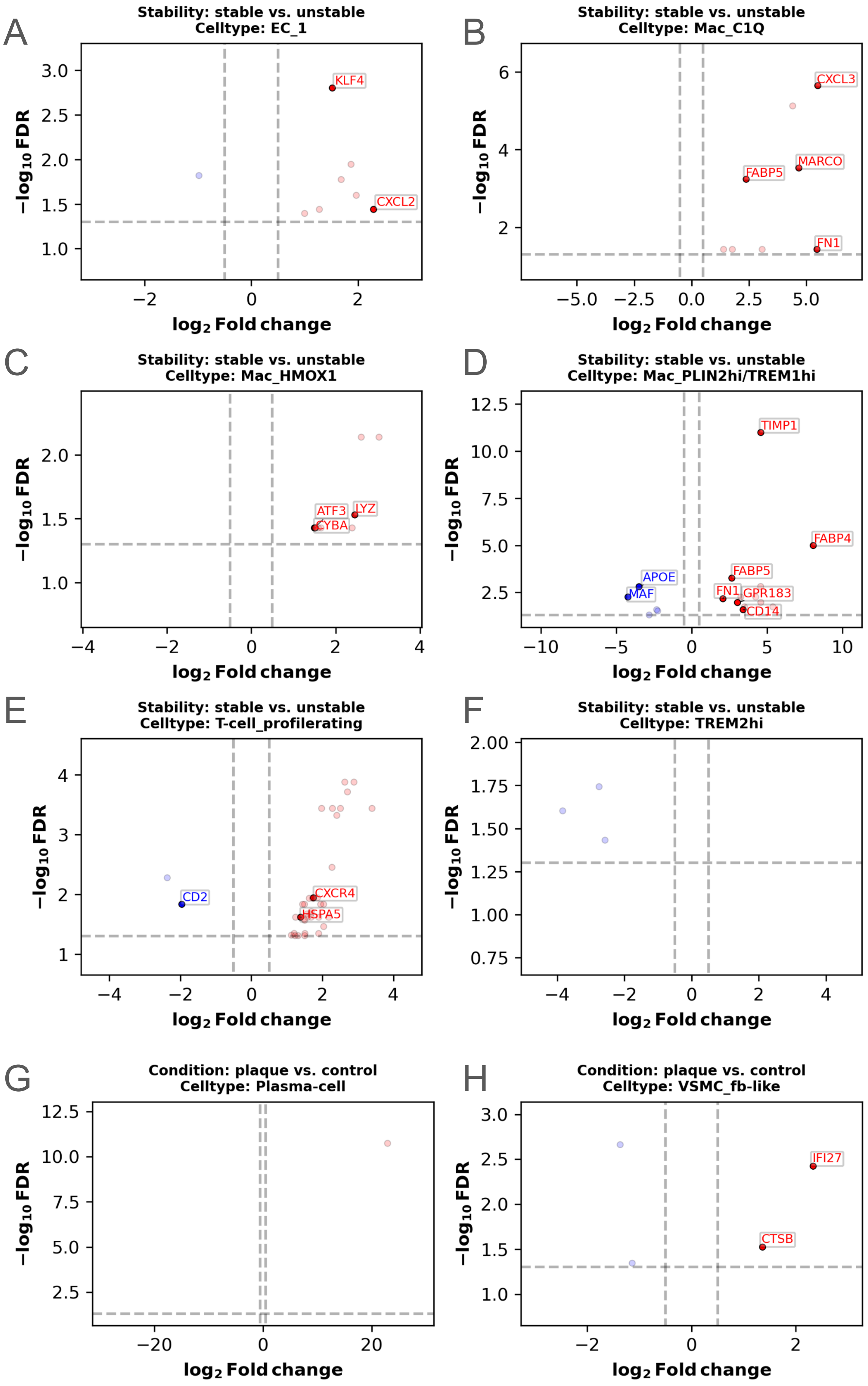


**Suppl. Fig. 1: Differential gene expression analysis results in the single-cell RNA-seq data.** Volcano plots depicting differentially expressed genes in the scRNA-seq dataset. Subplots (A - F) show the results of the condition comparison „stable vs. unstable“ within multiple low-level cell types respectively. (G - H): results of the condition comparison „plaque vs. control“ within multiple low-level cell types respectively. The x-axis refers to the $\log_2$ Fold change of the gene expression levels between the compared conditions, the y-axis shows the –log10 values of the Benjamini-Hochberg adjusted significance level (false discovery rate, FDR) of the gene expression level change. Vertical dashed line: log2 Fold change threshold at +/- 0.5, horizontal dashed line: FDR threshold of –log10(0.05). Red coloured genes are significantly up-, blue coloured genes are significantly downregulated in cells of the first condition in the comparison. Genes with name labels are also present in our Xenium gene panels.

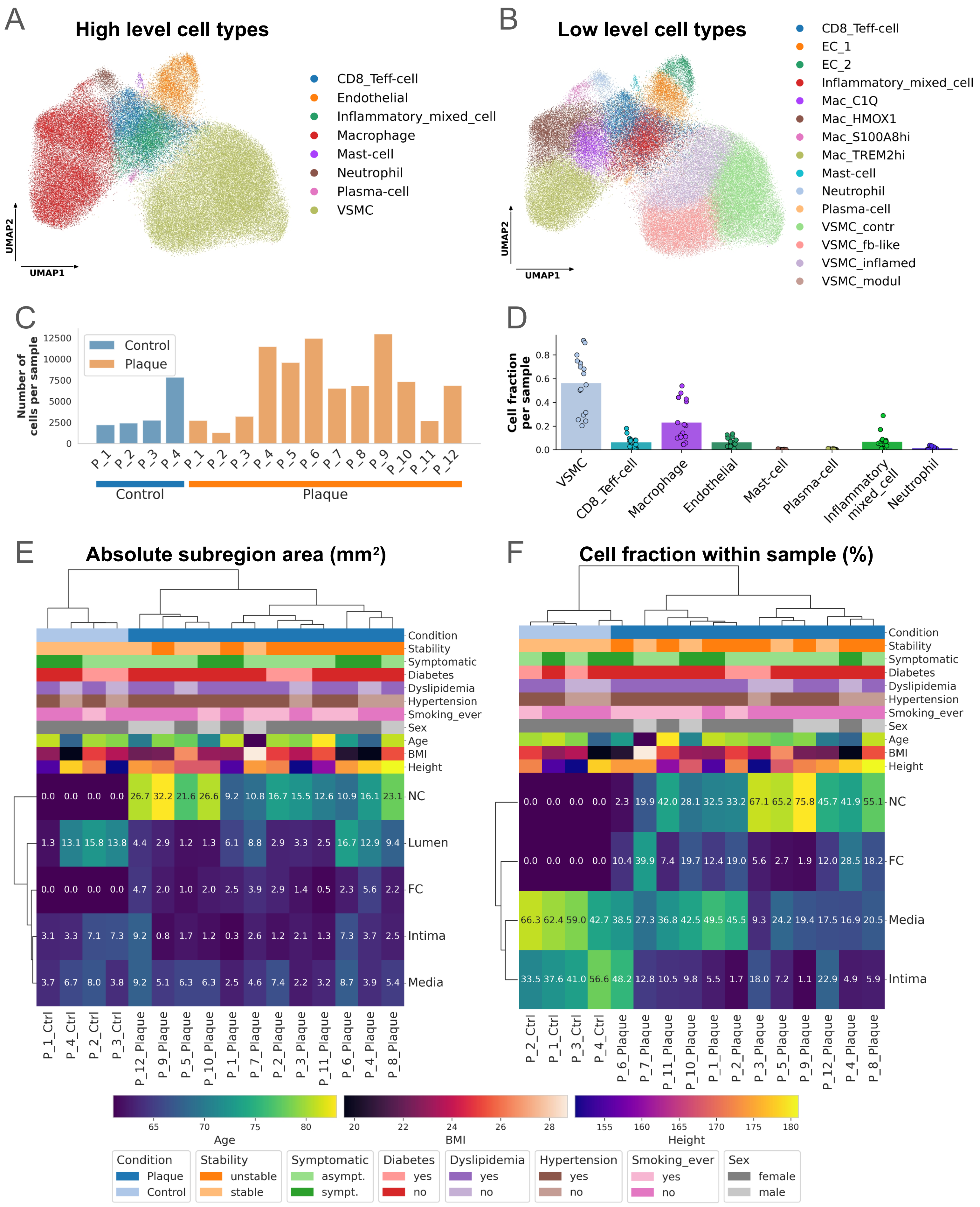


**Suppl. Fig. 2: Overview of cells coming from spatial transcriptomics with panel 2 genes and sample clusterings.** (A-B) UMAP low-dimensional representation of gene expression in cells of human atherosclerotic plaques, measured with Xenium method, panel 2 genes. Cells coloured by high-level cell types (A), cells coloured by low-level cell types (B). (C) Number of cells per sample, measured with Xenium panel 2 genes. (D) Cell fractions of high-level cell types per sample with panel 2 genes. (E) Hierarchical clustering of all samples based on absolute histological subregion areas. (F) Hierarchical clustering of all samples based on relative cell fractions in the 4 main cell-rich subregions. Cells were derived by Xenium panel 2 genes, normalisation was performed to total number of cells per sample, patient metadata are depicted above clustermap. „P“ stands for „Patient“ in the x-axis labels of (E) and (F), patient metadata legend is depicted below clustermaps (E) and (F). UMAP: Uniform Manifold Approximation and Projection, NC: necrotic core, FC: fibrous cap, sympt.: symptomatic, asympt.: asymptomatic.

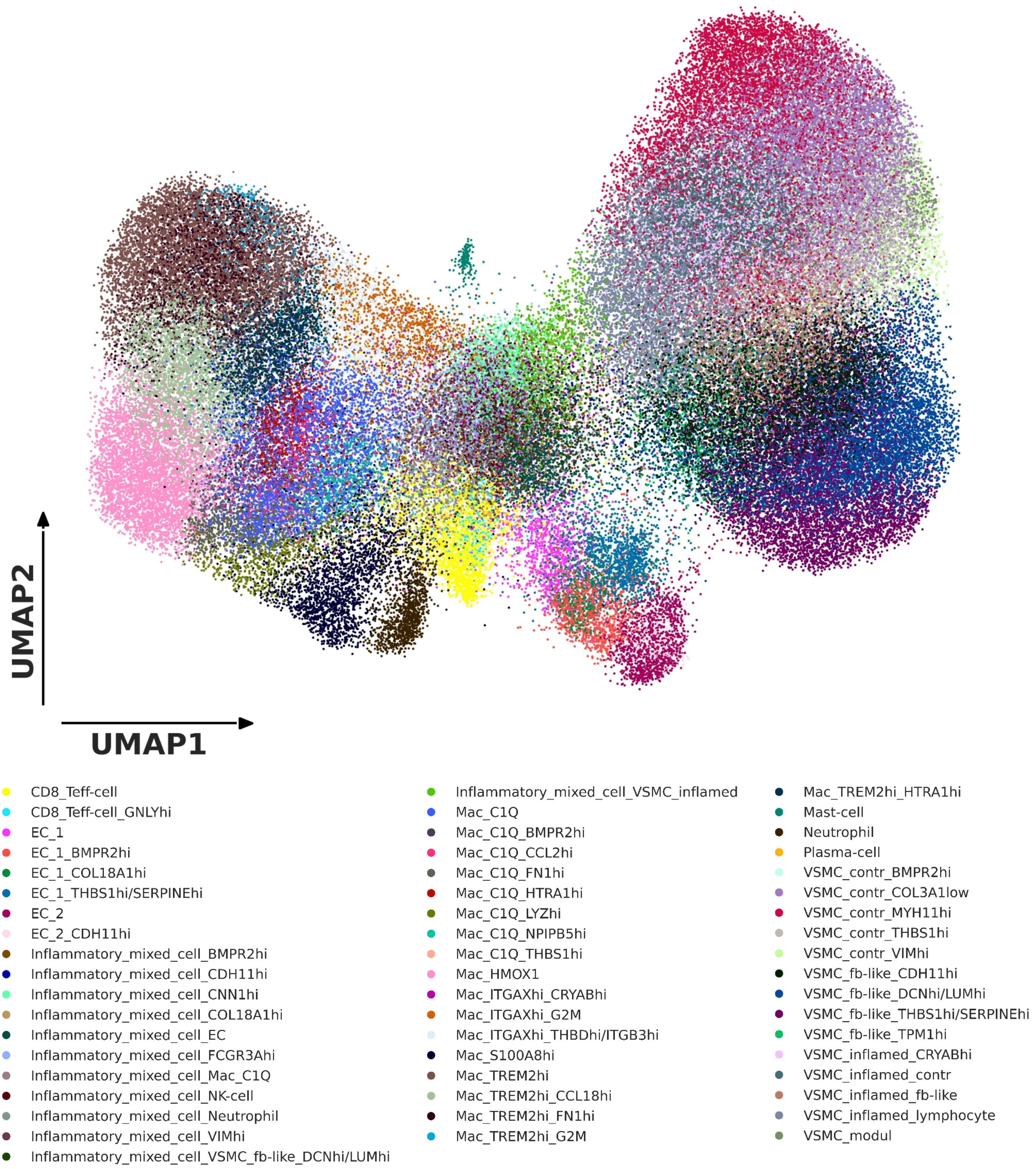


**Suppl. Fig. 3: UMAP showing the low-level cell subtype annotation of the final dataset with gene panel 1.** UMAP low dimensional representation of cell expression in the final dataset, gene panel 1. The colouring reflects the low-level cell substates annotation. UMAP: Uniform manifold maximum approximation and projection.

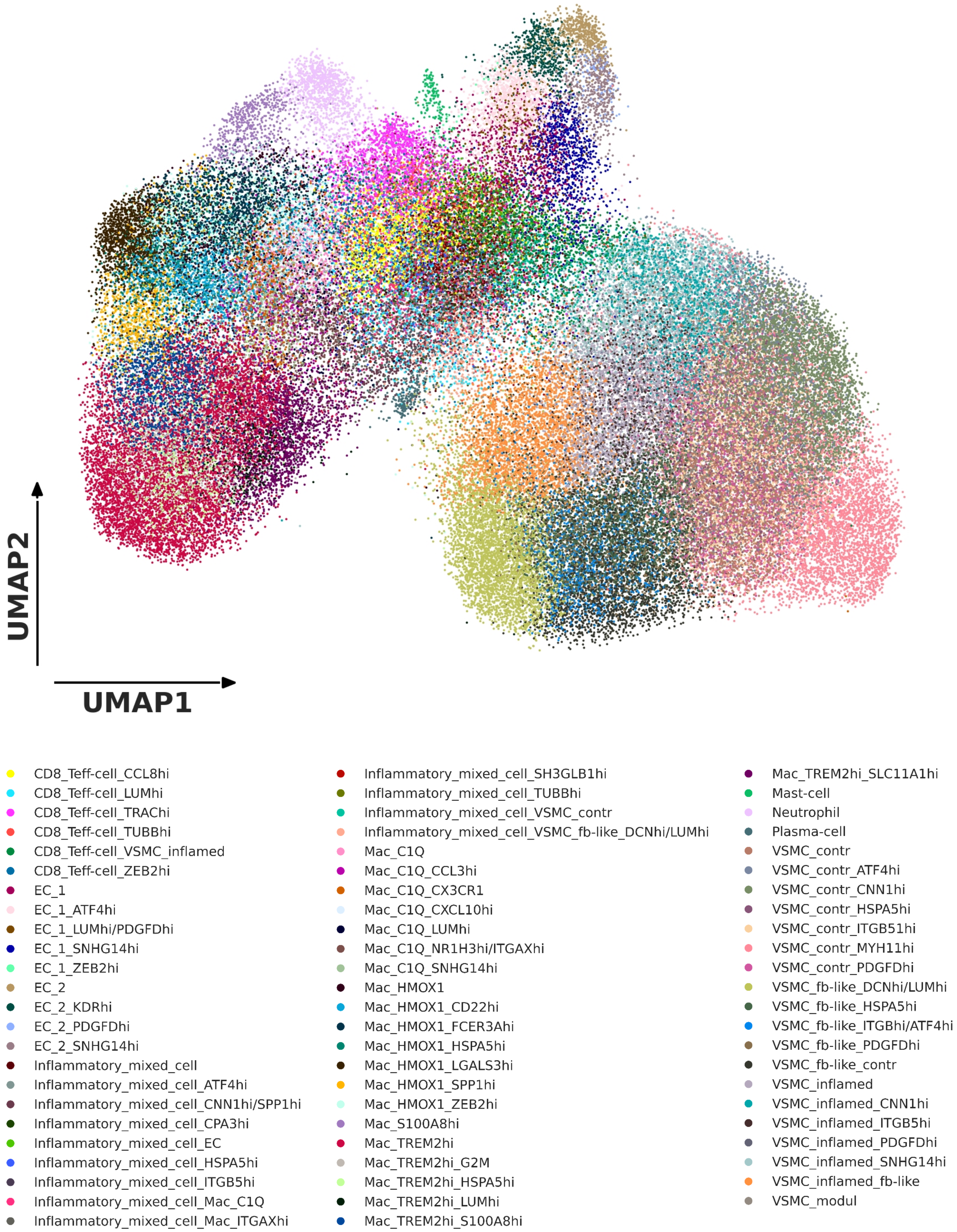


**Suppl. Fig. 4: : UMAP showing the low-level cell subtype annotation of the final dataset with gene panel 2.** UMAP low dimensional representation of cell expression in the final dataset, gene panel 2. The colouring reflects the low-level cell substates annotation. UMAP: Uniform manifold maximum approximation and projection

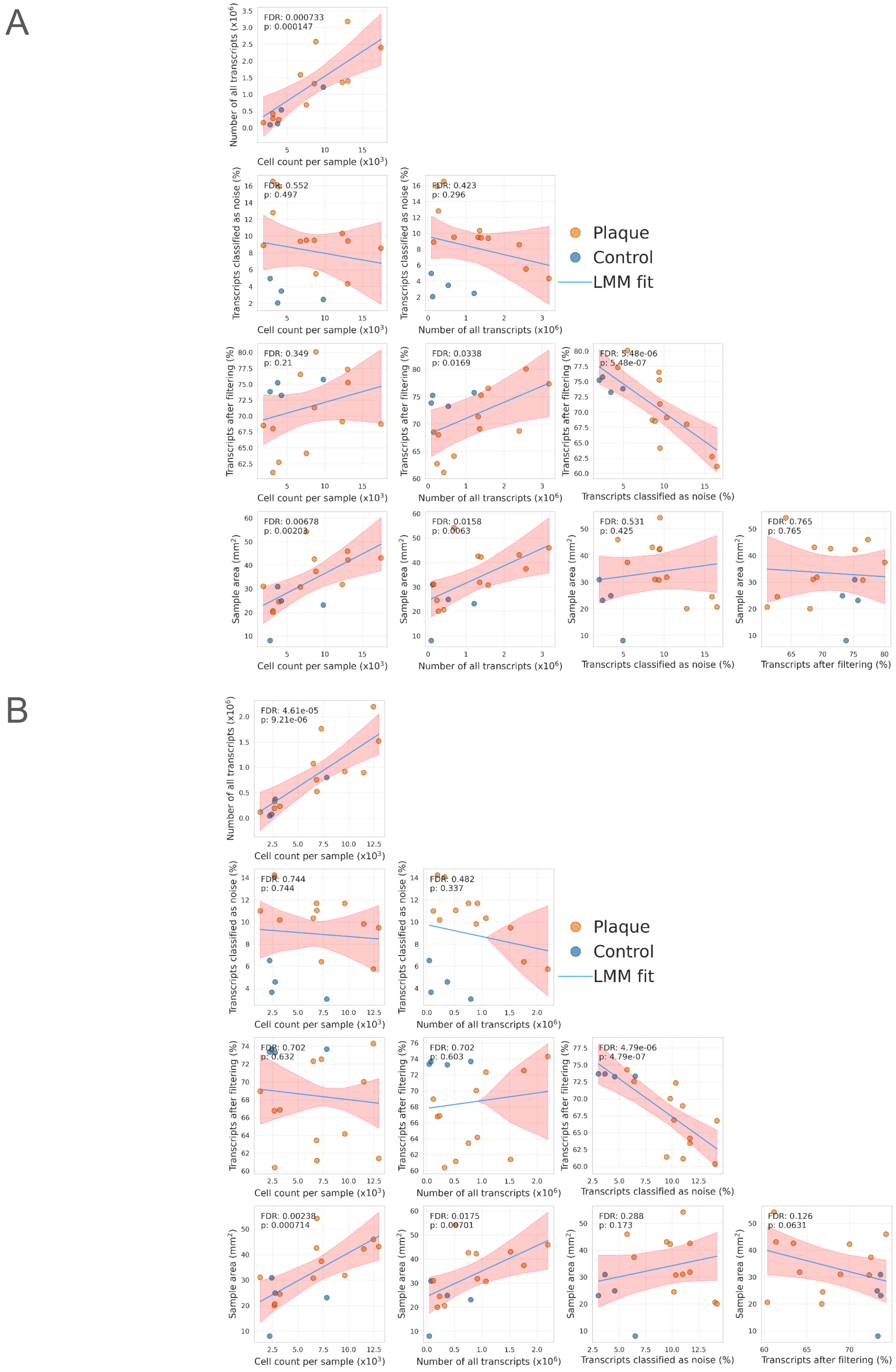


**Suppl. Fig. 7: Correlation between cell segmentation result metrics of final best cell segmentations per sample.** Correlations between cell segmentation result metrics of the final best cell segmentations selected for analysis for each sample with gene panel 1 (A), and gene panel 2 (B). As Patient 1-4 had Plaque and Control samples as well, correlation was measured by fitting a linear mixed model (LMM) adding random intercept effect for “Patient“. P-values represent the raw p-values of the LMM fit, false discovery rate (FDR) represents the Benjamini-Hochberg corrected raw LMM fit p-values for multiple testing.

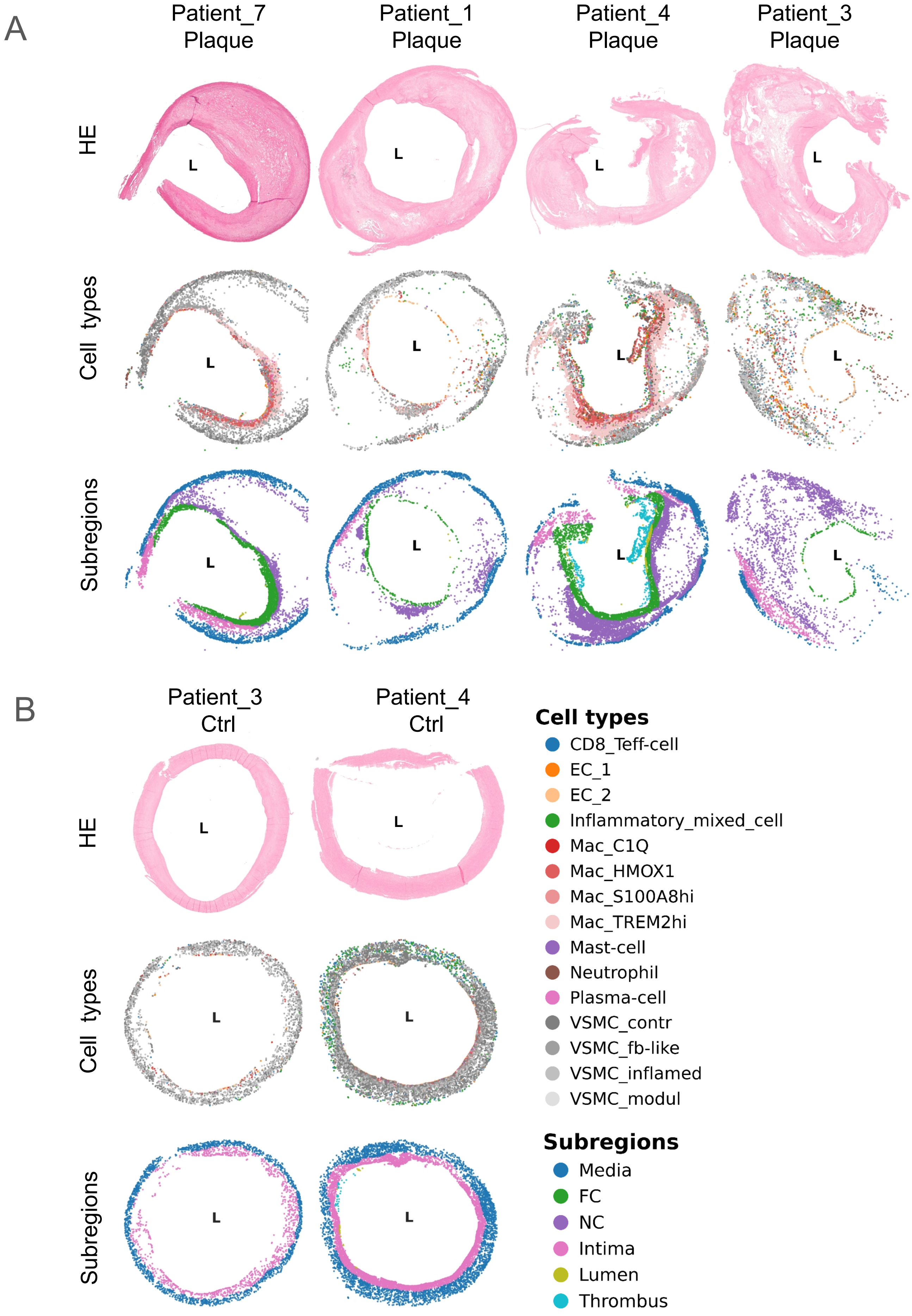


**Suppl. Fig. 8: General plaque compositions in panel 2.** (A) The upper row shows HE stainings of the plaques, the middle row indicates the the panel 2 low-level cell types, and the lower row shows the subregions of representative plaques in panel 2. L indicates the lumen. (B) The upper row shows HE stainings of the controls, the middle row indicates the the panel 2 low-level cell types, and the lower row shows the subregions of representative controls in panel 1. L indicates the lumen. HE: hematoxylin-eosin, NC: necrotic core, FC: fibrous cap.

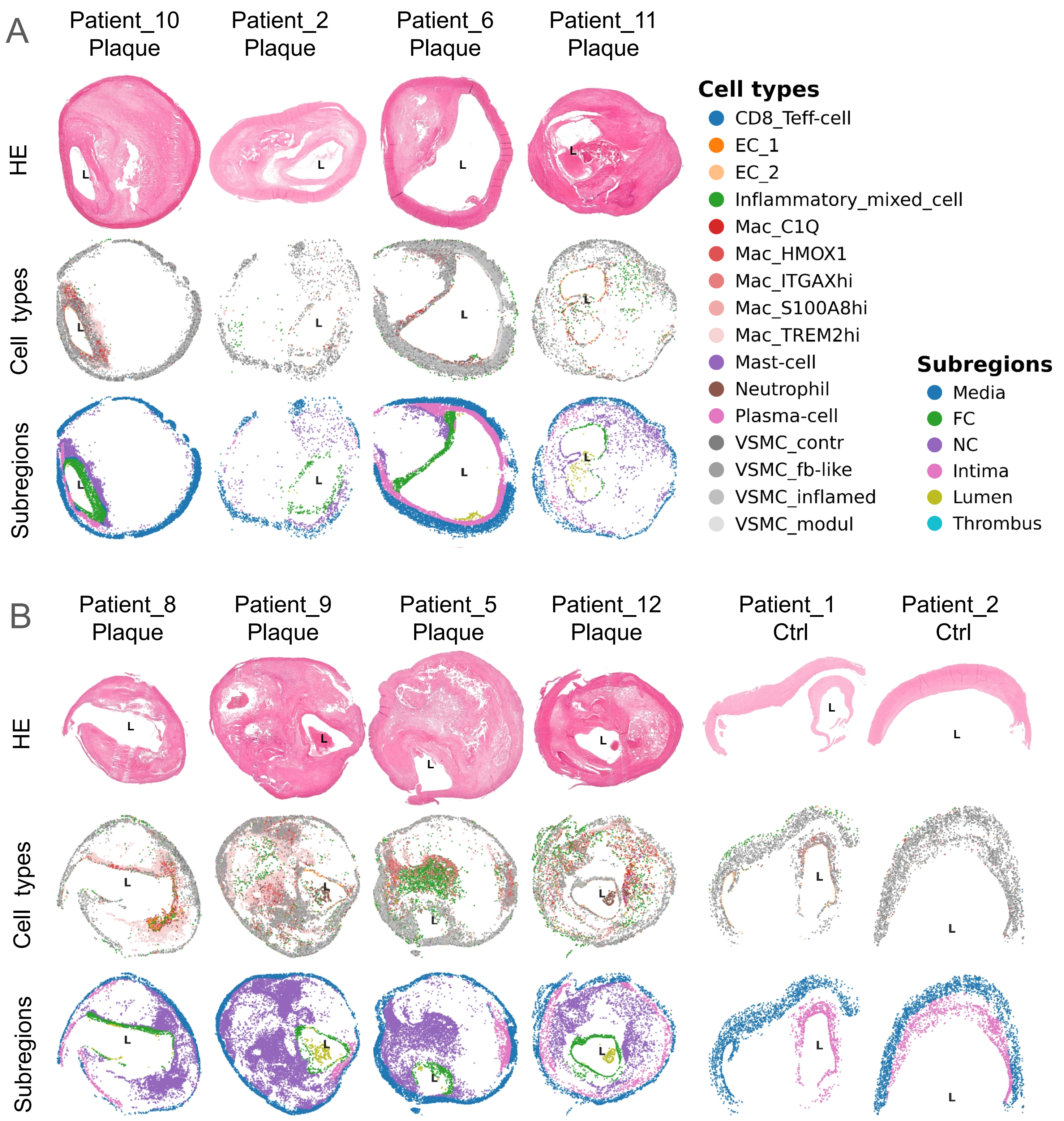


**Suppl. Fig. 9: General plaque compositions of all other plaques and controls in panel 1** (A) The upper row shows HE stainings of plaques, the middle row indicates the panel 1 low-level cell types, and the lower row shows the subregions of plaques in panel 1 (the rest of the plaques in shown in (B) or in Figure 3). L indicates the lumen. (B) The upper row shows HE stainings of plaques and control, the middle row indicates the panel 1 low-level cell types, and the lower row shows the subregions of plaques and controls in panel 1. L indicates the lumen. HE: hematoxylin-eosin, NC: necrotic core, FC: fibrous cap.

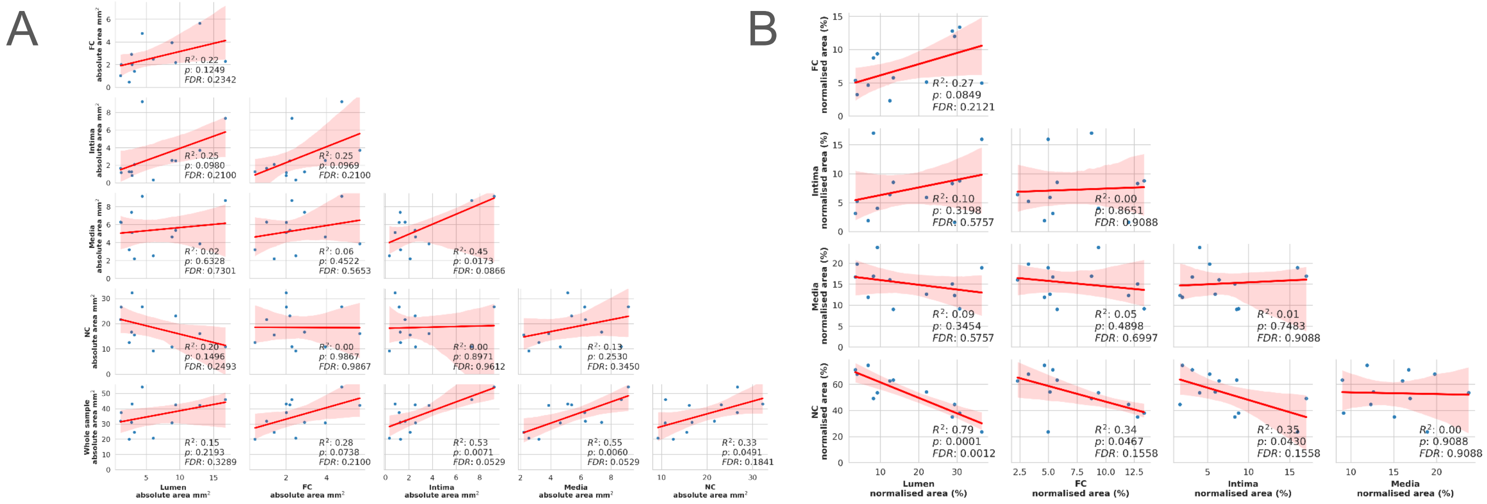


**Suppl. Fig. 10: Correlation of subregion and sample areas in plaques.** (A) Correlations between the absolute areas of individual subregions and the whole sample in our plaque samples. (B) Correlations between the normalised areas of individual subregions and in our plaque samples. R-squared and p-values come from a linear fit. Benjamini-Hochberg multiple testing correction was performed on all raw p-values within the absolute and normalised area correlations respectively and the corrected p-values are reported as false discovery rate (FDR). FC: fibrous cap, NC: necrotic core.

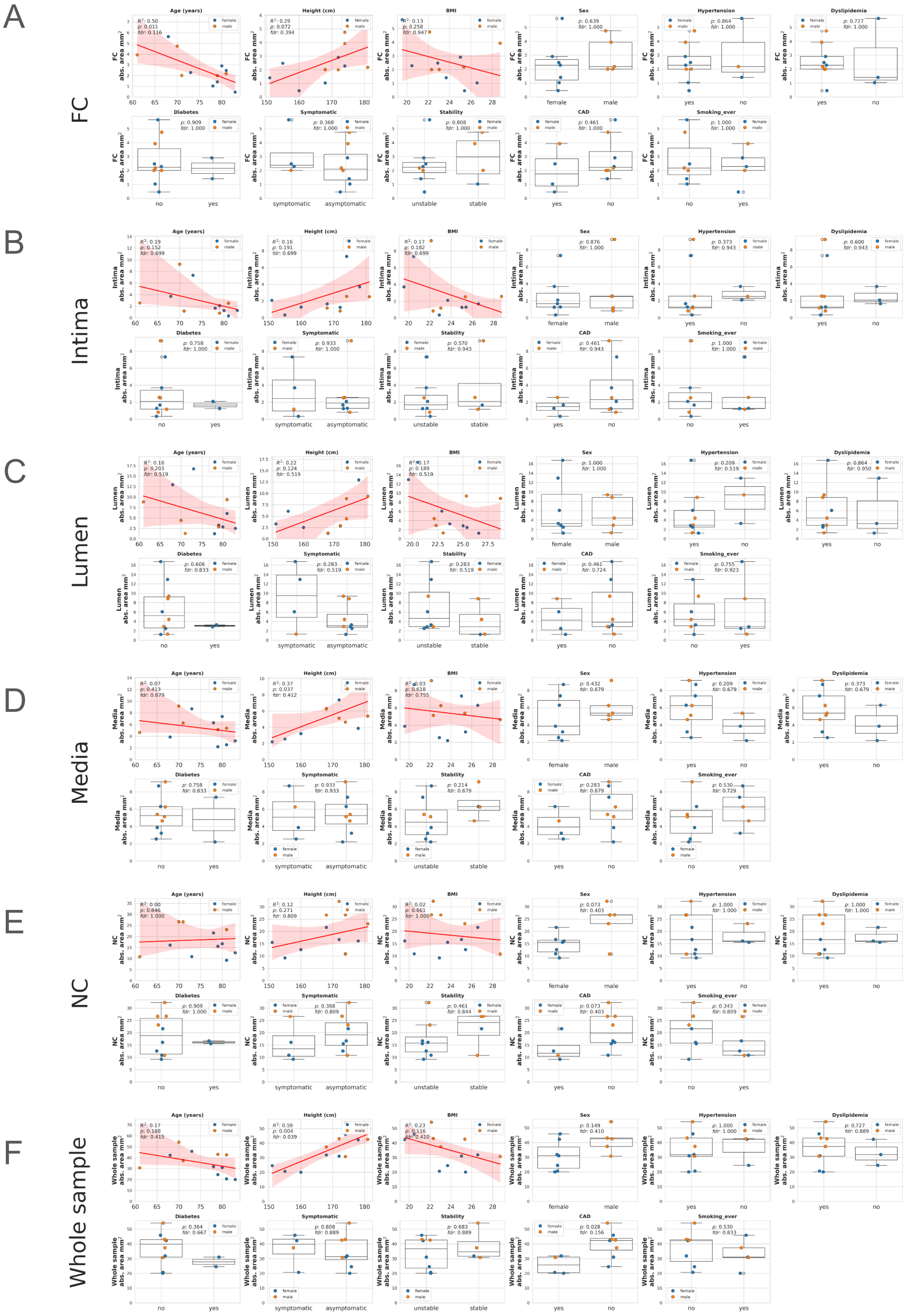


**Suppl. Fig. 11: Association of absolute subregion and sample areas of plaques with patient metadata, concomitant medical conditions and plaque features.** Association of absolute aubregion and sample areas of plaques with patient metadata, concomitant medical conditions and plaque features. Each subplot shows the associations of a different subregion: (A) fibrous cap (FC), (B) intima, (C) lumen, (D) media, (E) necrotic core (NC), (F) whole sample. For the numerical variables „Age“, „Height“ and „BMI“ R-squared and p-values come from a linear fit, for the remaining categorical variables Wilcoxon-tests were performed. Benjamini-Hochberg multiple testing correction was performed on all raw p-values within a subregion, the corrected p-values are reported as false discovery rate (FDR). COPD: chronic obstructive pulmonary disease, CAD: coronary artery disease.

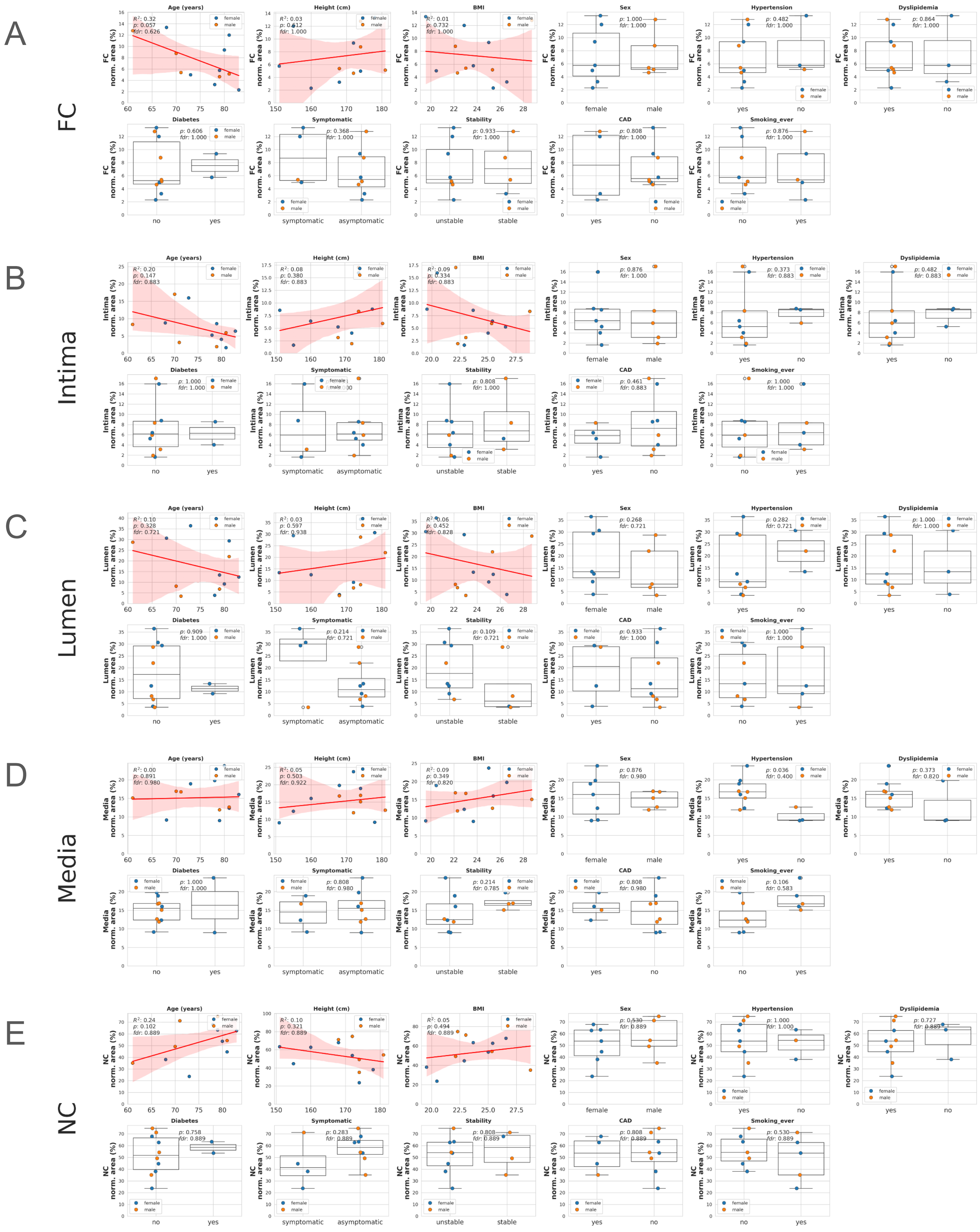


**Suppl. Fig. 12: Association of normalised subregion areas of plaques with patient metadata, concomitant medical conditions and plaque features.** Association of normalised subregion areas of plaques with patient metadata, concomitant medical conditions and plaque features. Each subplot shows the associations of a different subregion: (A) fibrous cap (FC), (B) intima, (C) lumen, (D) media, (E) necrotic core (NC). For the numerical variables „Age“, „Height“ and „BMI“ R-squared and p-values come from a linear fit, for the remaining categorical variables Wilcoxon-tests were performed. Benjamini-Hochberg multiple testing correction was performed on all raw p-values within a subregion, the corrected p-values are reported as false discovery rate (FDR). COPD: chronic obstructive pulmonary disease, CAD: coronary artery disease.

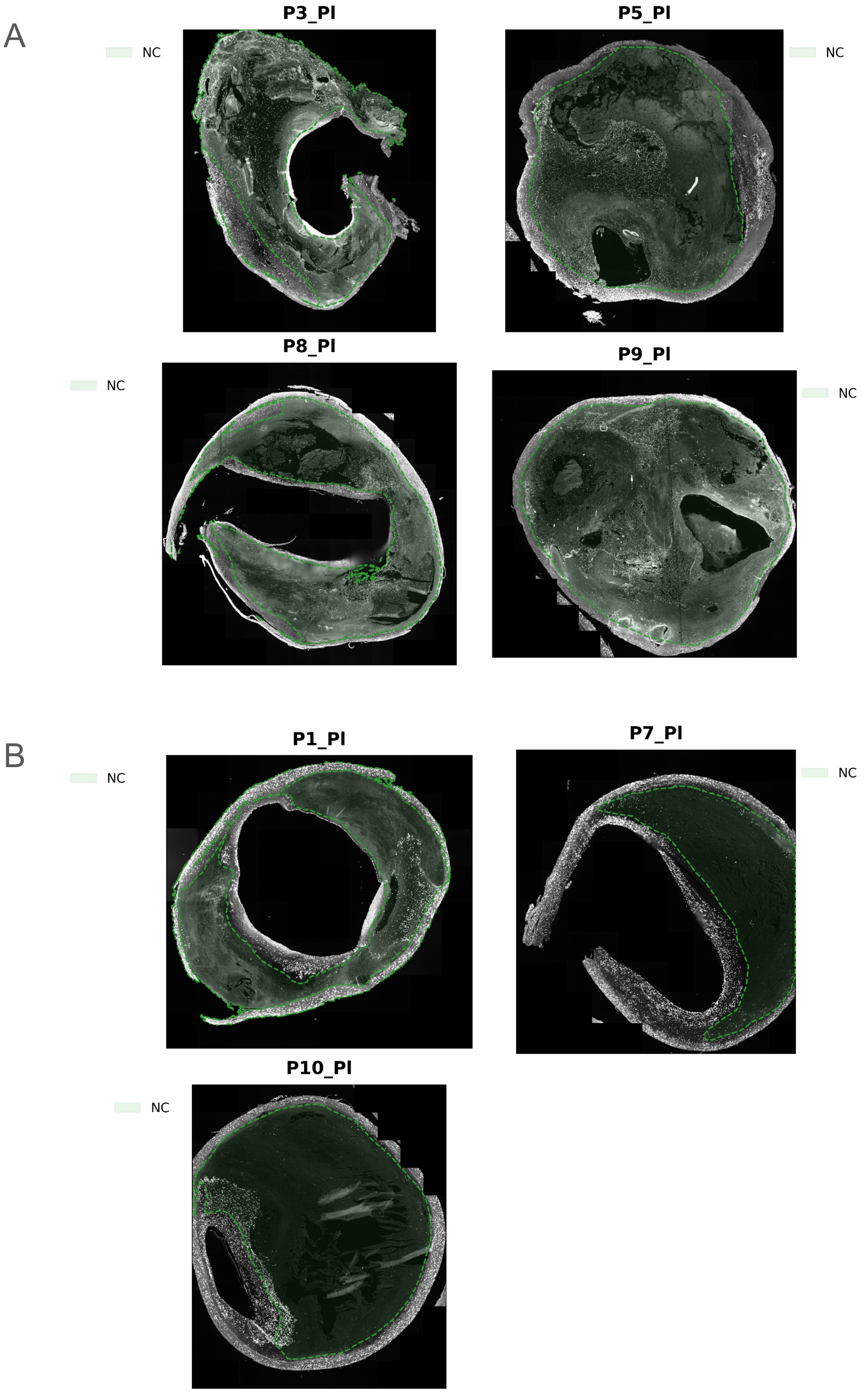


**Suppl. Fig. 13: DAPI-stained slides of plaques with high and low relative cell abundance in the necrotic core (NC).** (A) shows DAPI-stained slides of plaques (Patient 3,5,8,9) with large normalised necrotic core areas, that contain proportionally equally high fraction of cells. (B) shows DAPI-stained slides of plaques (Patient 1,7,10) with large normalised necrotic core areas, containing disproportionately low cell fractions compared to their normalised area. The necrotic core areas are coloured in a green shade.

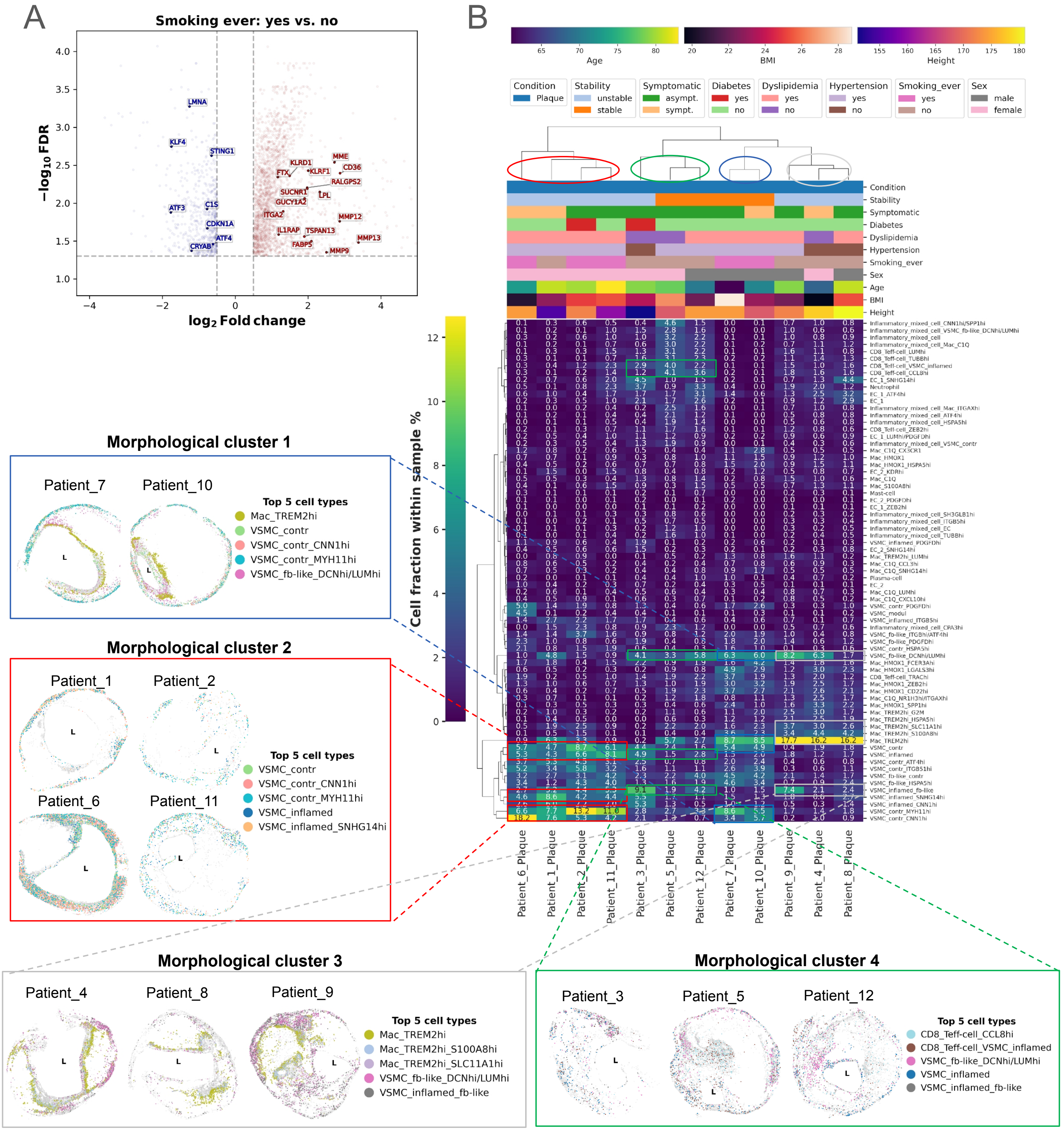


**Suppl. Fig. 14: Discordant differentially expressed genes across bulk and Xenium with "Smoking_ever" comparison, Morphological clusters in panel 2** (A) Bulk RNA-seq differential gene expression (DGE) results between „Smoking ever" conditions of plaques. The same DGE comparison was conducted with both Xenium gene panels. Labelled genes are significantly up or downregulated in both bulk and Xenium, with discordant log2 Fold change directions. FDR: false discovery rate, Benjamini-Hochberg adjusted significance level of gene expression change. Vertical dashed line: log2 Fold change threshold at +/- 0.5, horizontal dashed line: FDR threshold of –log10(0.05). (B) Heatmap shows hierarchical clustering of plaques into 4 morphological clusters based on panel 2 low-level substate cell fractions within each sample. Coloured boxes show spatial scatterplots of plaques grouped by morphological clusters. Dot colouring refers to the top 5 most abundant cell subtypes per cluster, „L" depicts lumen. FC: fibrous cap; NC: necrotic core; symp.: symptomatic, asymp.: asymptomatic.

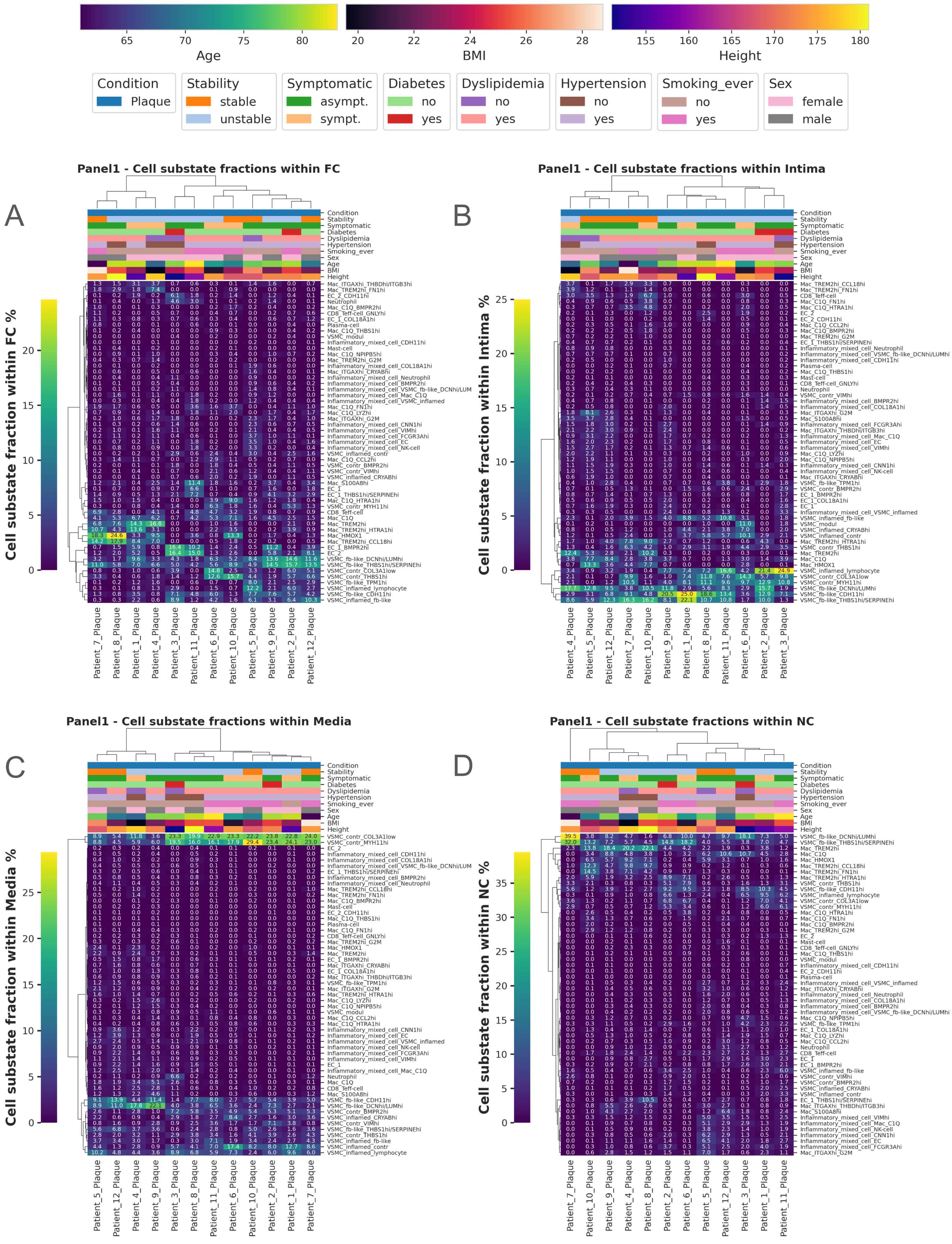


**Suppl. Fig. 15: Hierarchical clustering of plaque samples using the gene panel 1 low-level cell substate fractions normalised within different subregions per sample.** Subplots show the result of hierarchical clustering using the gene panel 1 low-level cell substate fractions normalised within (A) fibrous cap, (B) intima, (C) media, (D) necrotic core subregions. Column colouring depicts the distribution of patient metadata and plaque features across the clusters, with legend to the colouring placed at the top of the figure. The dendrogram above the column colouring represents the result of the clustering. FC: fibrous cap; NC: necrotic core; symp.: symptomatic, asymp.: asymptomatic.

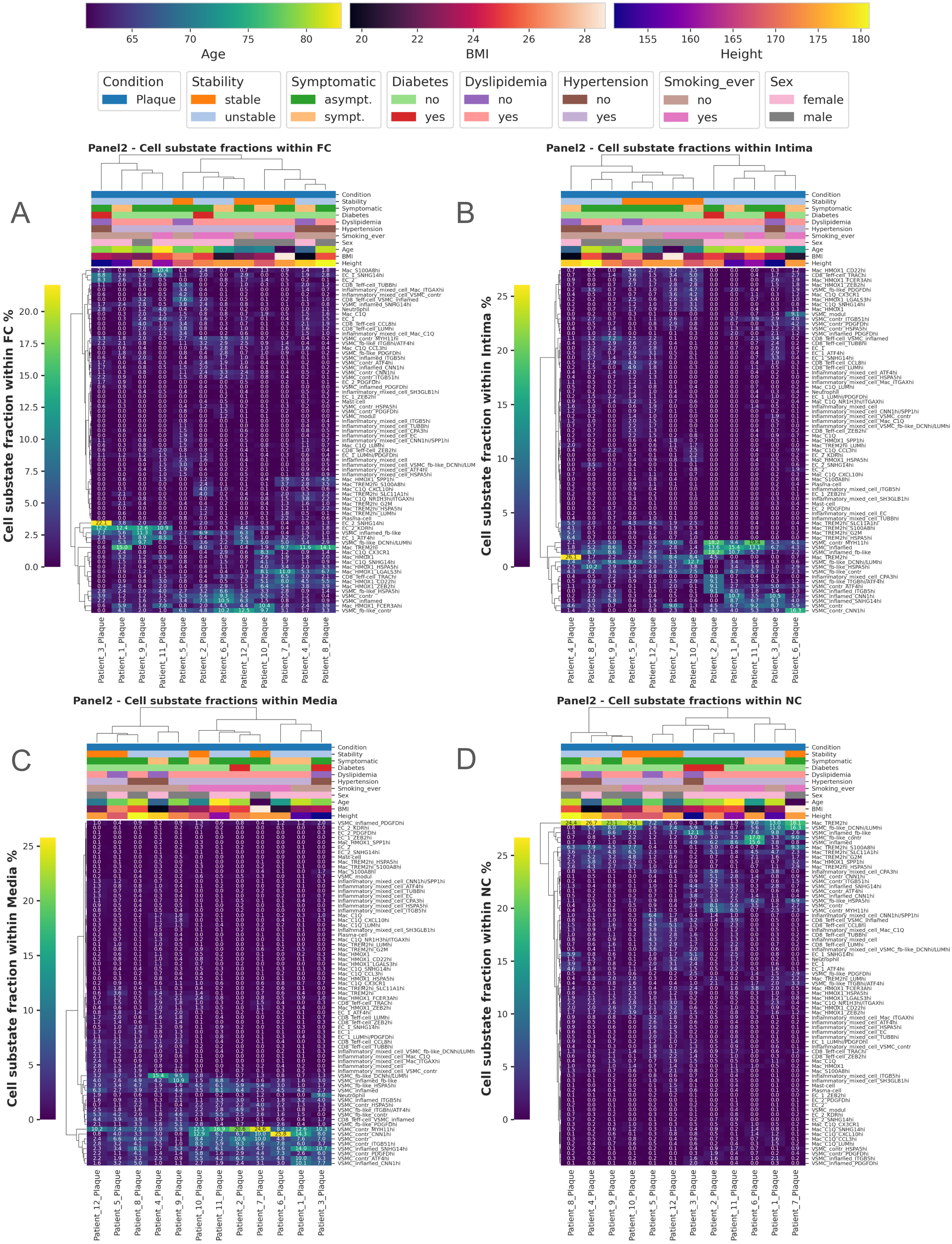


**Suppl. Fig. 16: Hierarchical clustering of plaque samples using the gene panel 2 low-level cell substate fractions normalised within different subregions per sample.** Subplots show the result of hierarchical clustering using the gene panel 2 low-level cell substate fractions normalised within (A) fibrous cap, (B) intima, (C) media, (D) necrotic core subregions. Column colouring depicts the distribution of patient metadata and plaque features across the clusters, with legend to the colouring placed at the top of the figure. The dendrogram above the column colouring represents the result of the clustering. FC: fibrous cap; NC: necrotic core; symp.: symptomatic, asymp.: asymptomatic

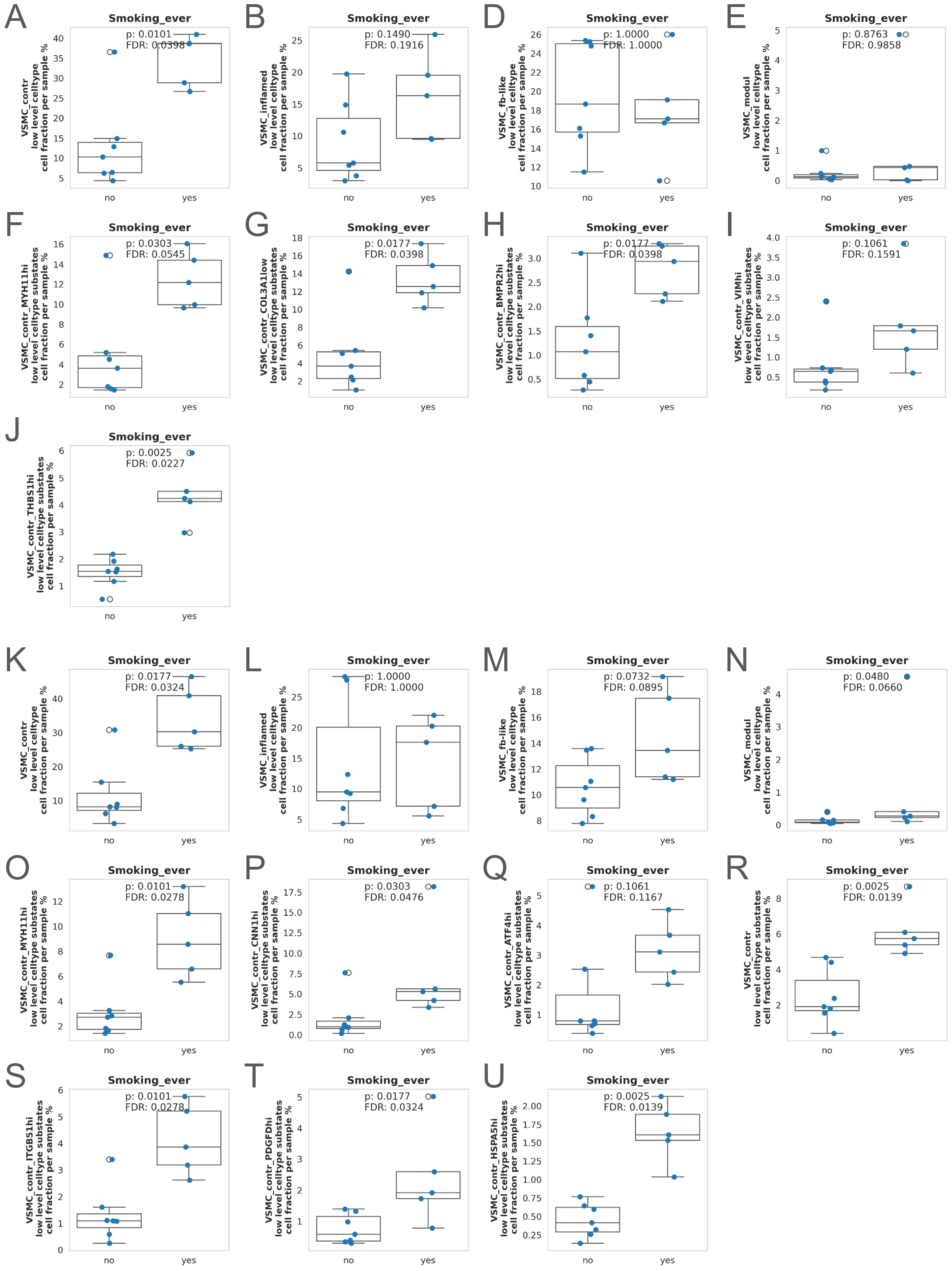


**Suppl. Fig. 17: Association of vascular smooth muscle cell (VSMC) subtype cell fractions in Xenium plaques with „Smoking ever" status.** Boxplots showing the association of different VSMC subtype fractions of plaque samples with Smoking ever" status of the patients (no=7, yes=5). The y-axis label describes the cell subtype name and the annotation level it is from (low-level cell type / low-level cell substate). (A-E) show the cell fractions of the 4 VSMC low-level cell types of gene panel 1, (F-J) show the contractile VSMC substate fractions of gene panel 1, (K-N) show the cell fractions of the 4 VSMC low-level cell types of gene panel 2, (F-J) show the contractile VSMC substate fractions of gene panel 2. P-values refer to Wilcoxon-tests performed across „Smoking ever" status levels. Benjamini-Hochberg multiple testing correction was performed on all raw p-values, the corrected p-values are reported as false discovery rate (FDR).

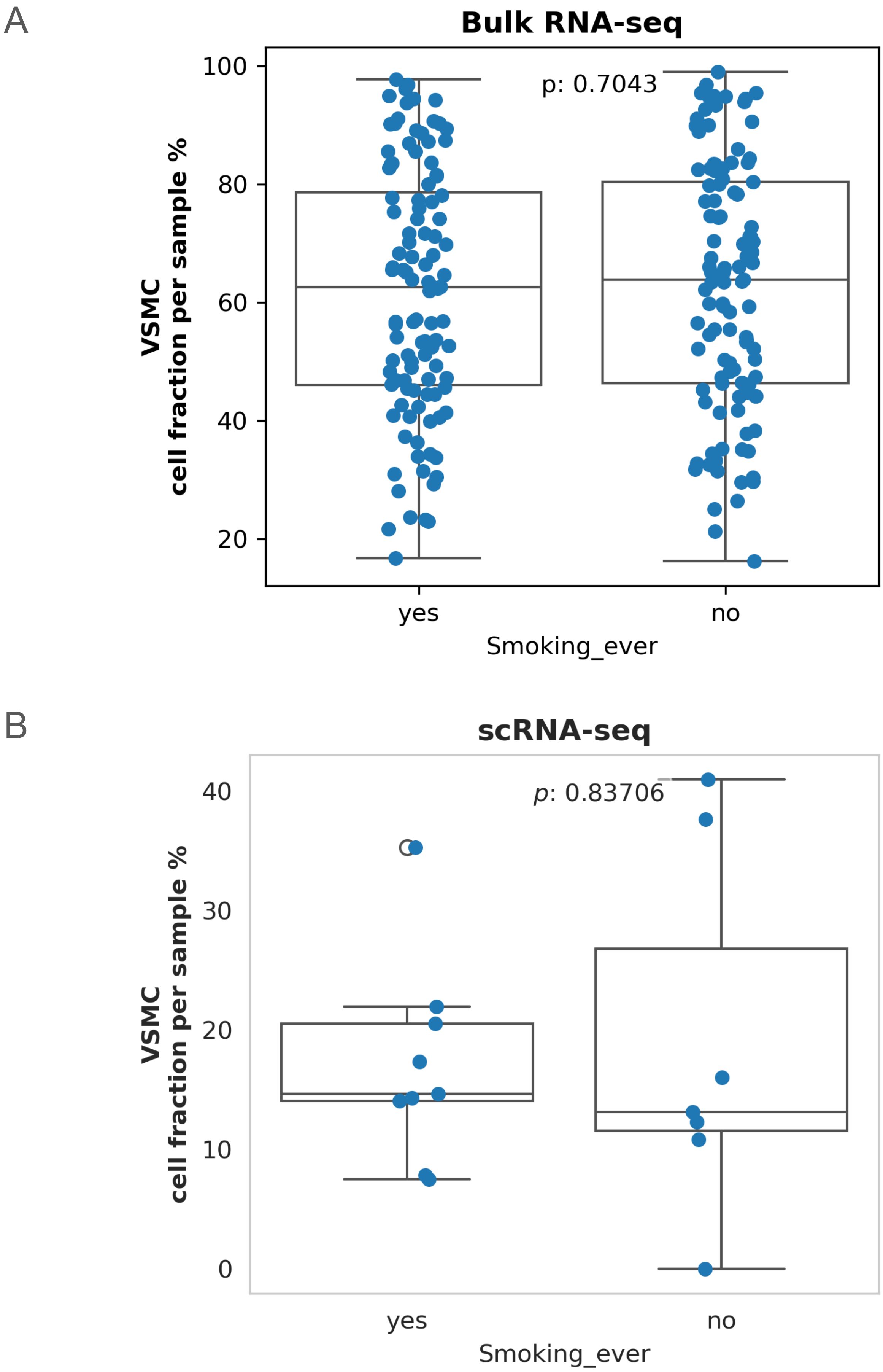


**Suppl. Fig. 18: Association of vascular smooth muscle cell (VSMC) subtype cell fractions in plaques with „Smoking ever“ status.** Boxplots showing the association of VSMC high-level cell type fractions of plaque samples with Smoking ever“ status of the patients. (A) VSMC cell fractions per Smoking_ever label (no=109, yes=108) in our bulk RNA-seq data. Cell fractions derived from deconvolution. (B) VSMC cell fractions per Smoking_ever label (no=7, yes=9) in our scRNA-seq data. P-values represent significance level of the performed Wilcoxon-test.

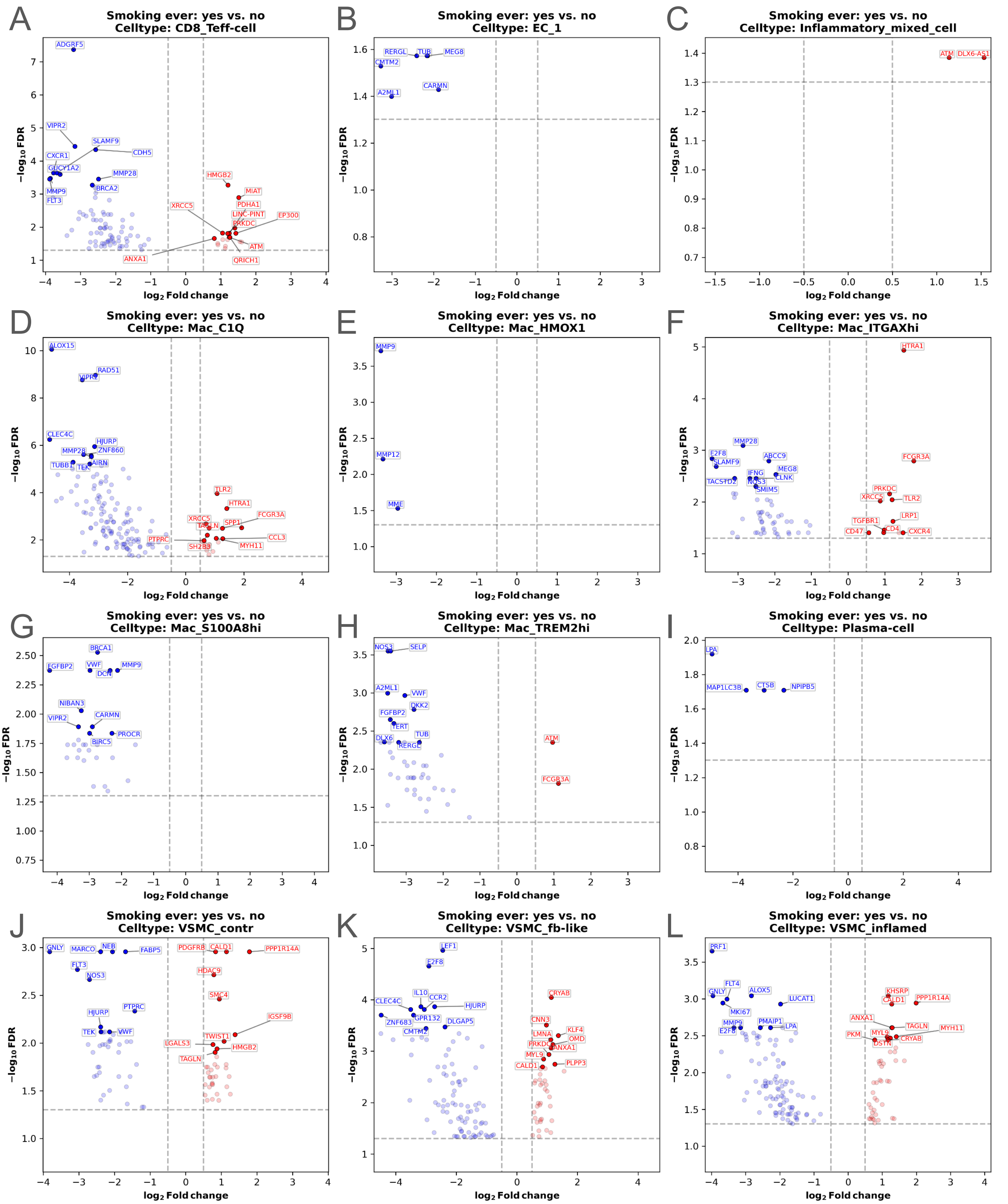


**Suppl. Fig. 19: Differential gene expression result of „Smoking ever" comparison of plaques with Xenium panel 1 genes.** Volcano plots showing the results of differential gene expression (DGE) analysis performed across „Smoking ever" conditions in plaques with Xenium panel 1 gene dataset. (A-L) DGE analysis was performed within low-level cell types. X-axis depicts the log2 fold-change of gene expression across „Smoking ever" conditions, y-axis depicts the –log10 values of the Benjamini-Hochberg adjusted significance level of gene expression change (false discovery rate, FDR). Gene name labels represent the top 10 most significantly upregulated (red) and downregulated (blue) genes. Vertical dashed lines: log2 Fold change threshold at +/- 0.5, horizontal dashed line: FDR threshold of –log10(0.05).

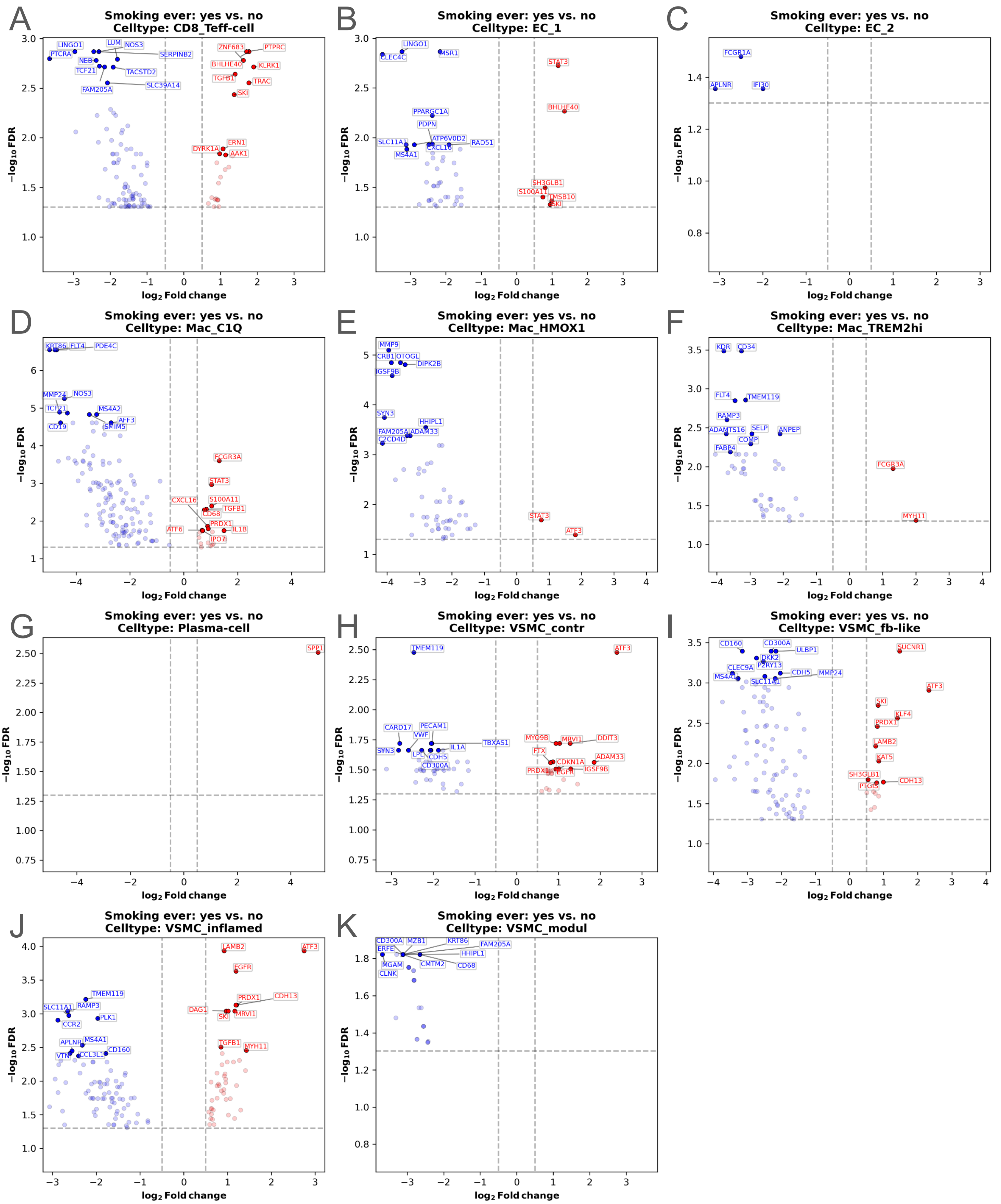


**Suppl. Fig. 20: Differential gene expression result of „Smoking ever" comparison of plaques with Xenium panel 2 genes.** Volcano plots showing the results of differential gene expression (DGE) analysis performed across „Smoking ever" conditions in plaques with Xenium panel 2 gene dataset. (A-K) DGE analysis was performed within low-level cell types. X-axis depicts the log2 fold-change of gene expression across „Smoking ever" conditions, y-axis depicts the –log10 values of the Benjamini-Hochberg adjusted significance level of gene expression change (false discovery rate, FDR). Gene name labels represent the top 10 most significantly upregulated (red) and downregulated (blue) genes. Vertical dashed lines: log2 Fold change threshold at +/- 0.5, horizontal dashed line: FDR threshold of –log10(0.05).

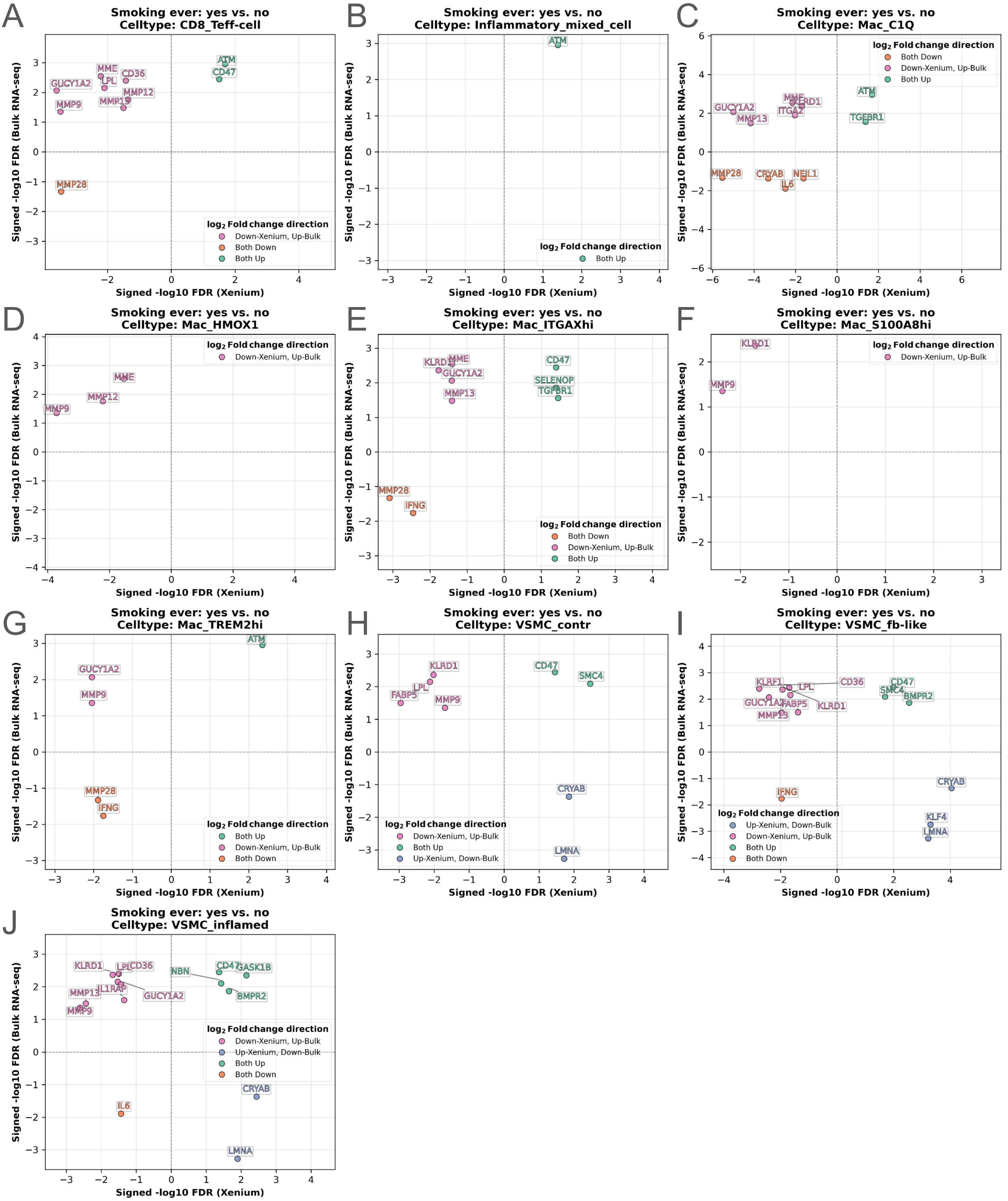


**Suppl. Fig. 21: Signed FDR plots comparing differential gene expression (DGE) results between Xenium panel 1 data and bulk RNA-seq in plaque samples**. Differential expression analysis was performed separately in each low-level cell type within the Xenium panel 1 dataset, while the bulk RNA-seq analysis was conducted across all plaque-derived samples. Each subplot (A-J) corresponds to a specific low-level cell type from Xenium, showing genes that were significantly differentially expressed in both datasets based on the "Smoking ever" condition. The x-axis shows the signed -log10 FDR values from Xenium gene panel 1, while the y-axis represents the signed -log10 FDR values from bulk RNA-seq. Dot colouring represents the log2 fold change relations of genes in both datasets. FDR: false discovery rate, Benjamini-Hochberg adjusted significance level of gene expression change.

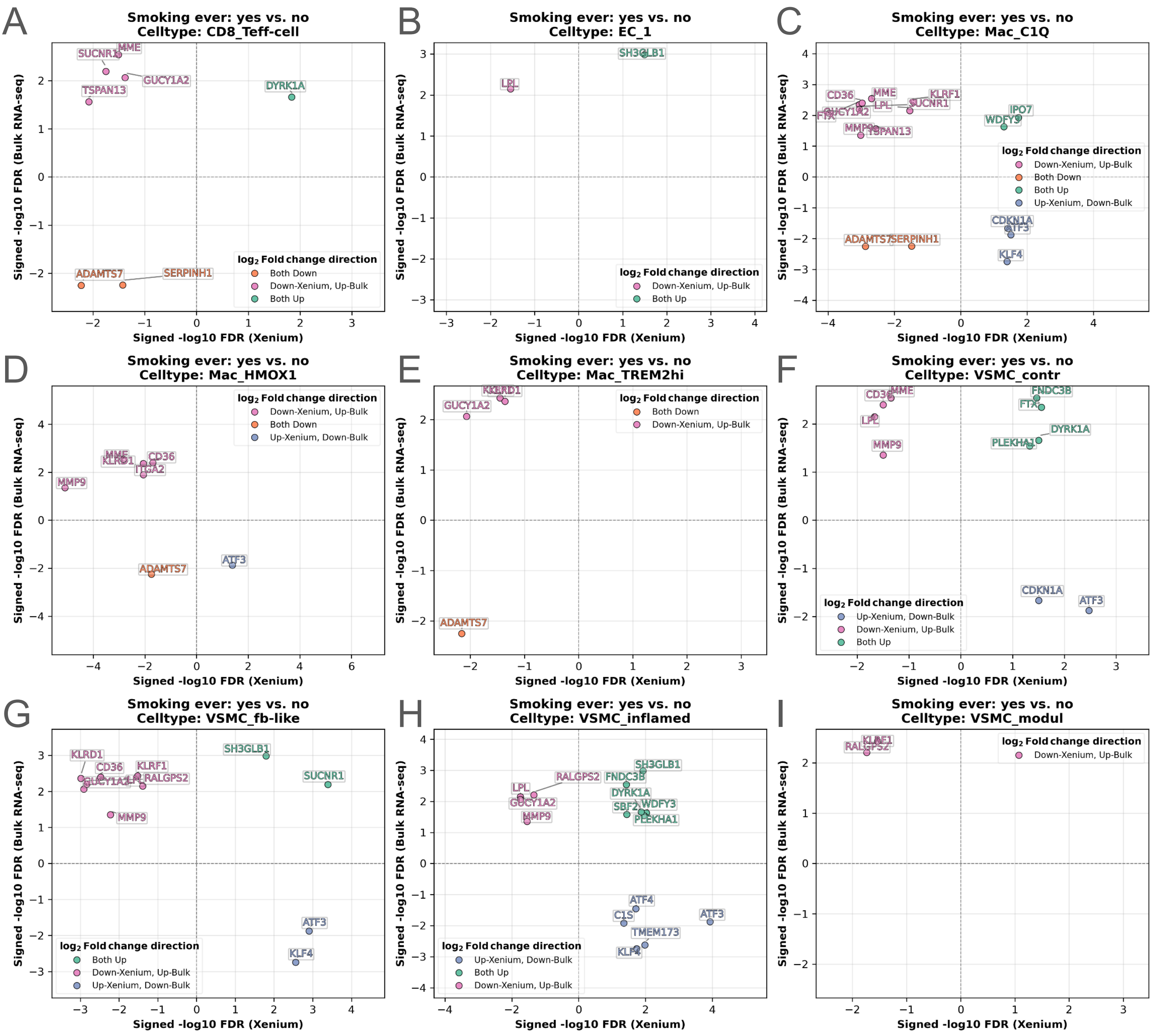


**Suppl. Fig. 22: Signed FDR plots comparing differential gene expression (DGE) results between Xenium panel 2 data and bulk RNA-seq in plaque samples**. Differential expression analysis was performed separately in each low-level cell type within the Xenium panel 2 dataset, while the bulk RNA-seq analysis was conducted across all plaque-derived samples. Each subplot (A-I) corresponds to a specific low-level cell type from Xenium, showing genes that were significantly differentially expressed in both datasets based on the "Smoking ever" condition. The x-axis shows the signed -log10 FDR values from Xenium gene panel 1, while the y-axis represents the signed -log10 FDR values from bulk RNA-seq. Dot colouring represents the log2 fold change relations of genes in both datasets. FDR: false discovery rate, Benjamini-Hochberg adjusted significance level of gene expression change.

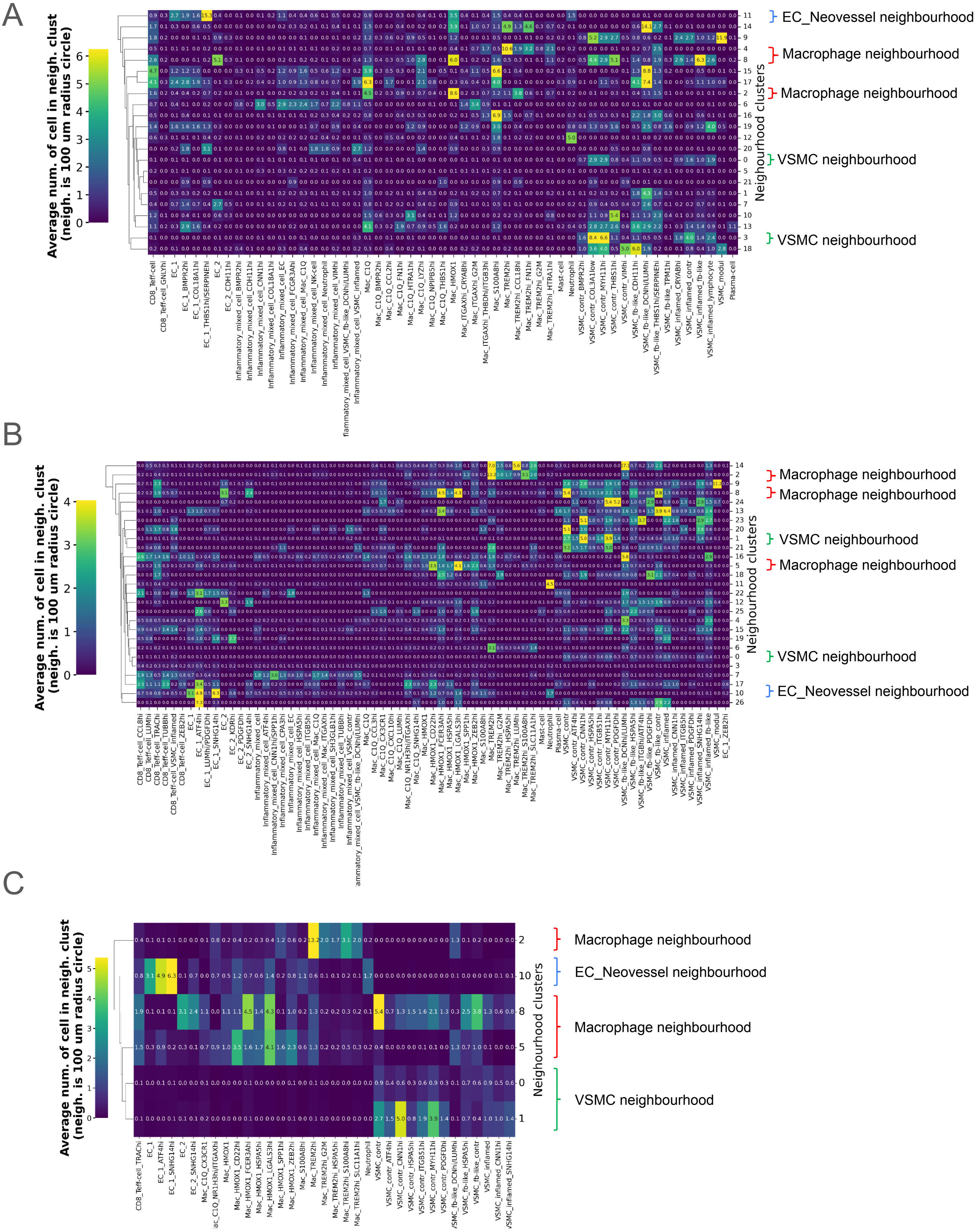


**Suppl. Fig. 23: All cell neighbourhoods in panel 1 and panel 2.** Clustermaps showing the average local neighbourhoods (signatures) of all panel 1 (A) and all panel 2 (B) neighbourhood clusters. Local neighbourhood of a cell is defined as the number of neighbouring cell substates within a 100 um radius (for more details see Methods and Results). Vascular smooth muscle cell (VSMC) neighbourhoods of panel 1: cluster 0 & 3, panel 2: cluster 0 & 1. macrophage neighbourhoods of panel 1: cluster 2, 4 & 8, panel 2: 2,5 & 8. Endothelial cell (EC) neovessel neighbourhood of panel 1: cluster 1, panel 2: 11. (C) Clustermap showing the average local neighbourhoods of the panel 2 VSMC, Macrophage and EC neovessel neighbourhood clusters. For brevity, only the most abundant substates are shown on the x-axis.

A

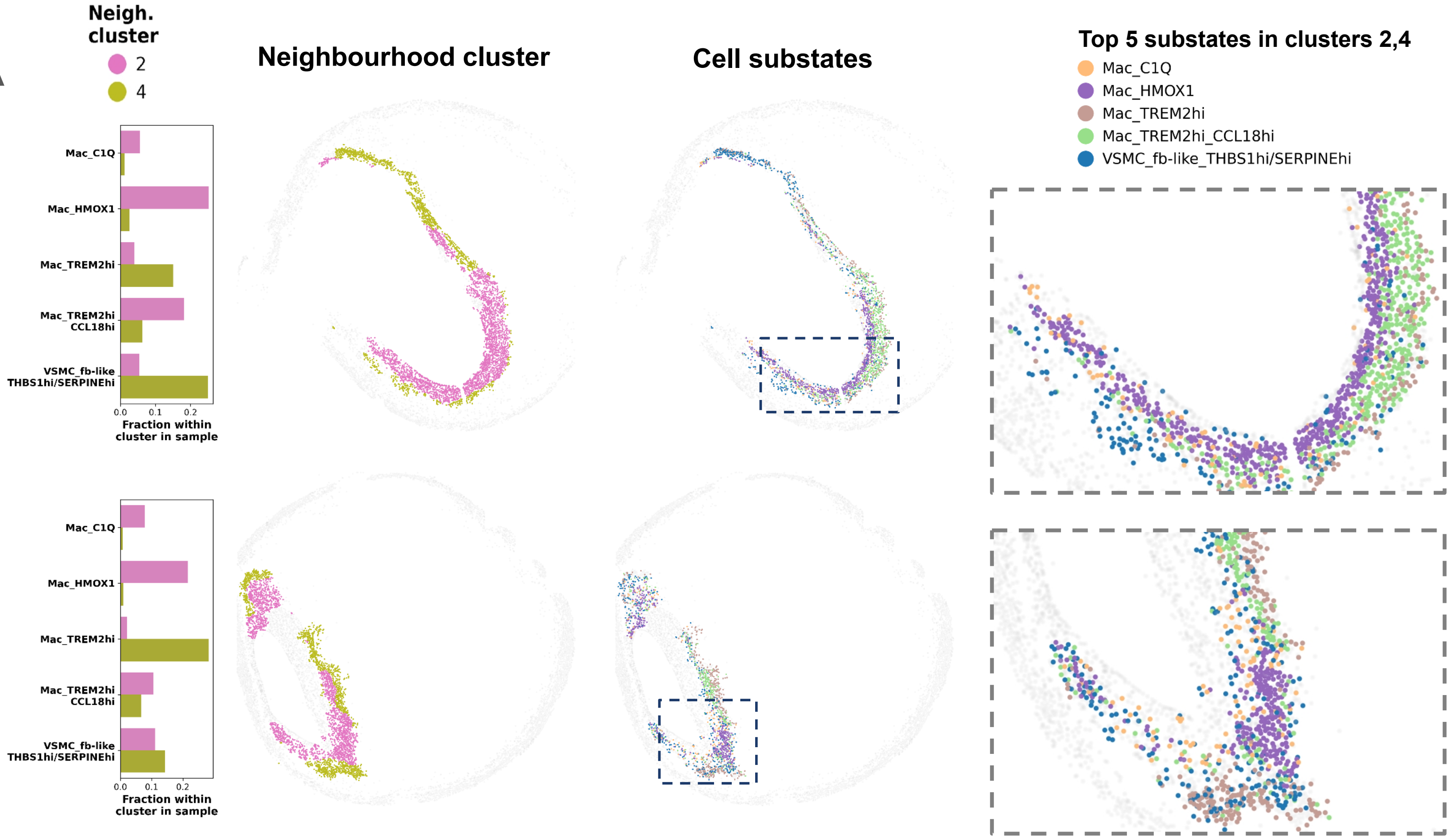


B

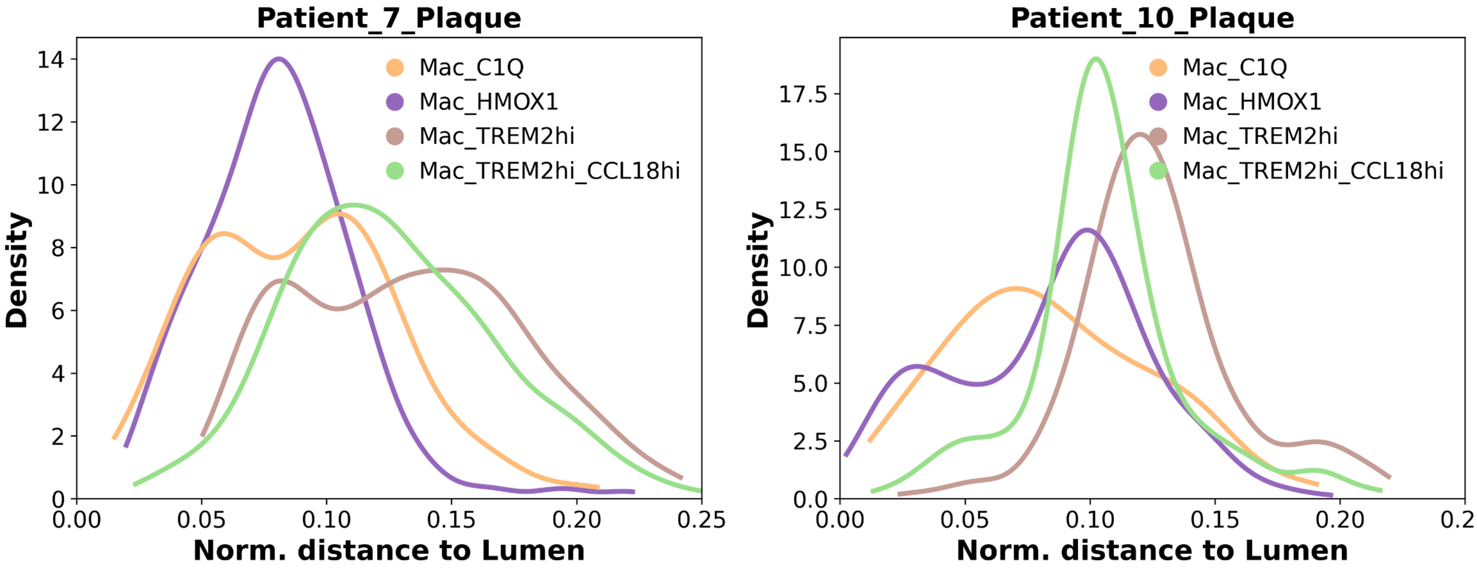


**Suppl. Fig. 24: Macrophage neighbourhoods presented in further plaques.** Composition of macrophage neighbourhoods shown based on the five most abundant cell substates per panel 1 neighbourhood cluster, in two further plaques. The first column displays relative fractions of these substates within each neighbourhood cluster per plaque. The second column shows their spatial distribution coloured by neighbourhood cluster, while the third column presents the same distribution coloured by cell substate. The fourth column provides a zoomed-in view of regions with interesting substate spatial distribution. Dashed rectangles in column three represent the original location of interesting regions within sample. (B) Density plot showcasing the ordered layering of four macrophage substates in the two selected plaques. X-axis depicts the normalised distance from lumen, (capped at 0.25 to only consider subluminal regions), y-axis depicts the distribution of distances.

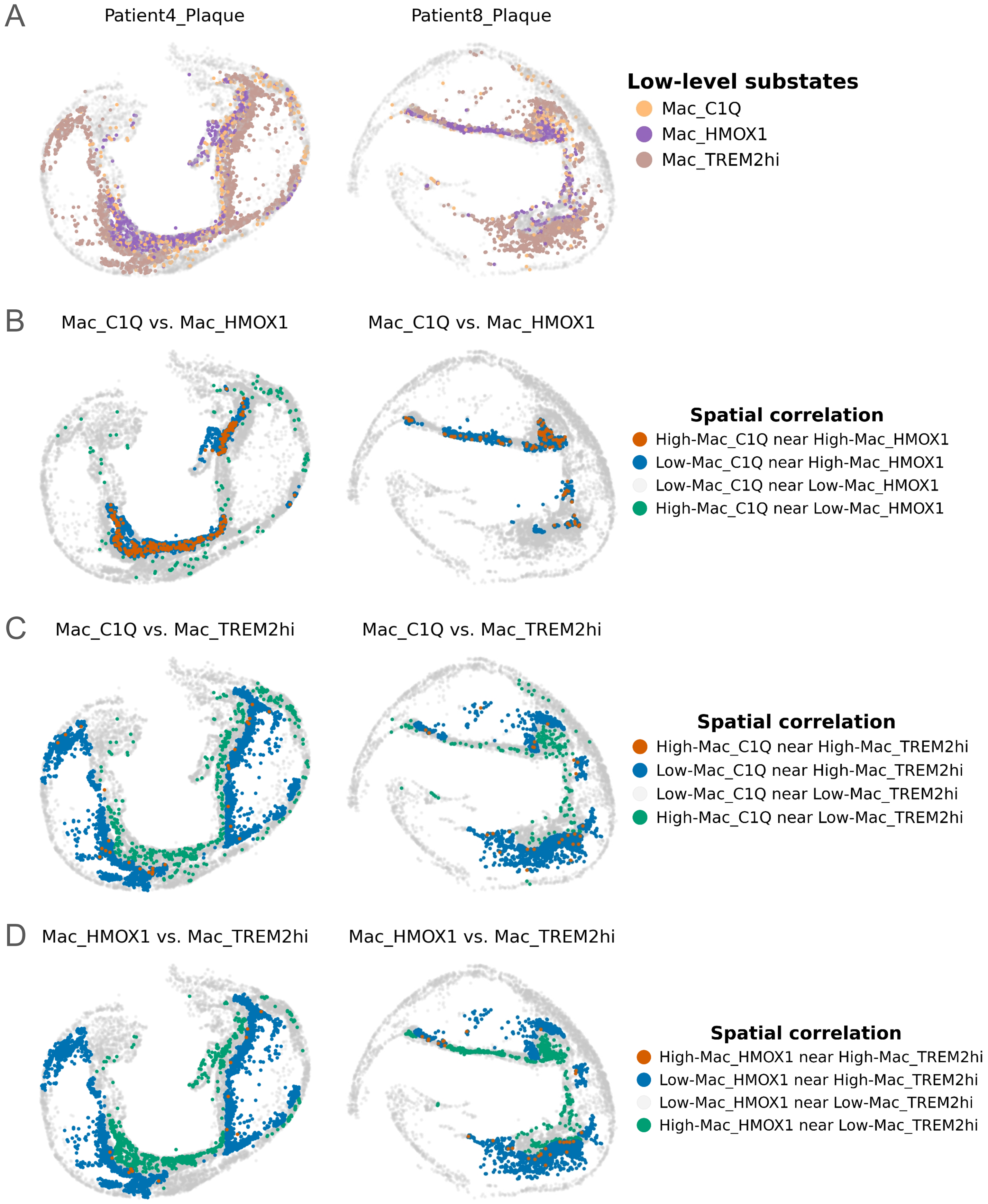


**Suppl. Fig. 25: Spatial correlation of Macrophage substates.** (A) Spatial distribution of the panel 1 Macrophage substates Mac_C1Q, Mac_HMOX1 and Mac_TREM2hi for two selected samples. (B-D) Spatial correlation calculated between substate pair combinations using Local Bivariate Moran's I statistic. Cells coloured other than grey show a significant spatial correlation across substate pairs (FDR<0.05), indicating the nature of correlation (i.e. high-high, high-low). FDR: false discovery rate, resulting from Benjamini-Hochberg correction of raw p-values of Local Bivariate Moran's I values. Multiple testing correction was performed within sample per substate pair.

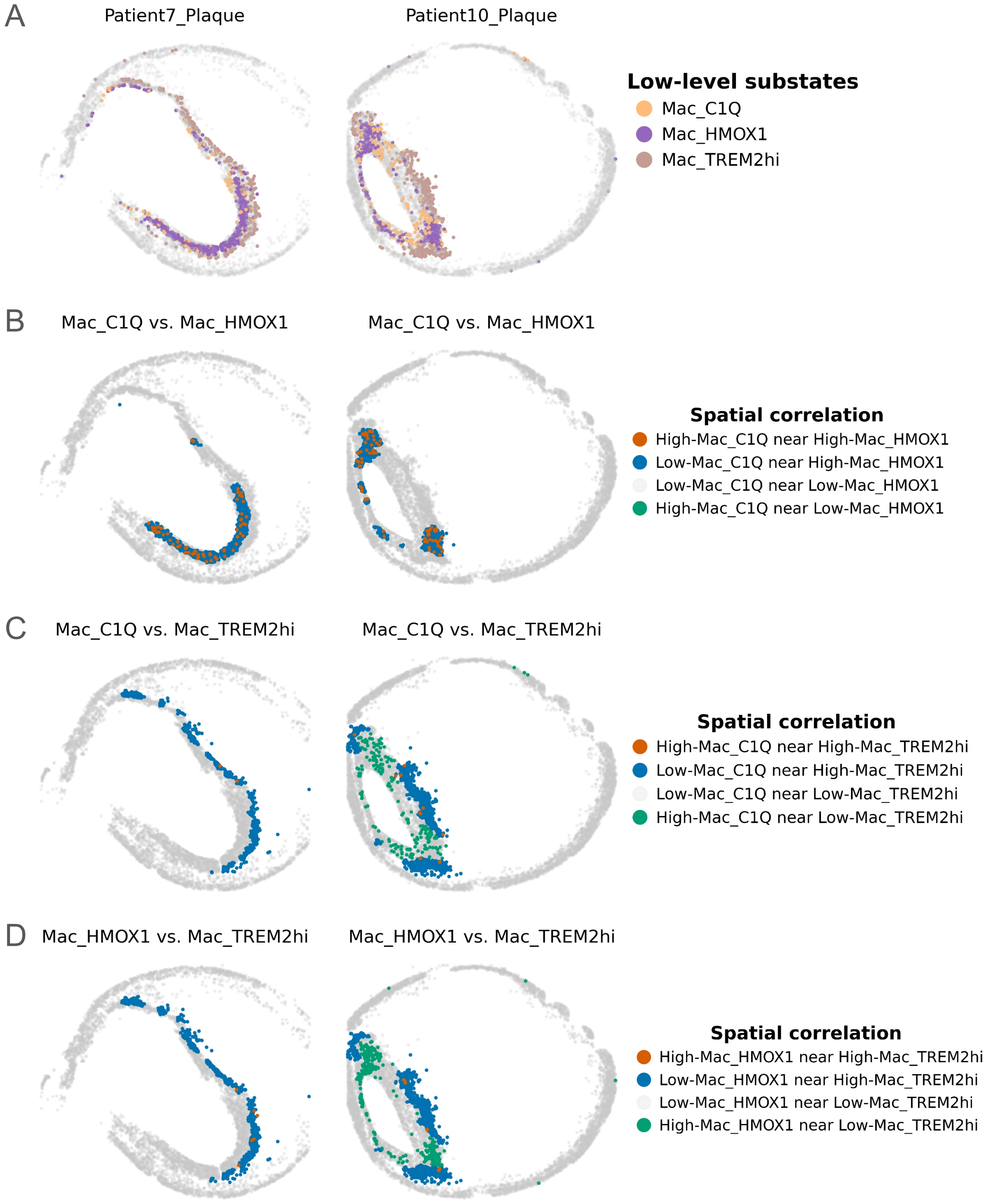


**Suppl. Fig. 26: Spatial correlation of Macrophage substates.** (A) Spatial distribution of the panel 1 Macrophage substates Mac_C1Q, Mac_HMOX1 and Mac_TREM2hi for two selected samples. (B-D) Spatial correlation calculated between substate pair combinations using Local Bivariate Moran's I statistic. Cells coloured other than grey show a significant spatial correlation across substate pairs (FDR<0.05), indicating the nature of correlation (i.e. high-high, high-low). FDR: false discovery rate, resulting from Benjamini-Hochberg correction of raw p-values of Local Bivariate Moran's I values. Multiple testing correction was performed within sample per substate pair.

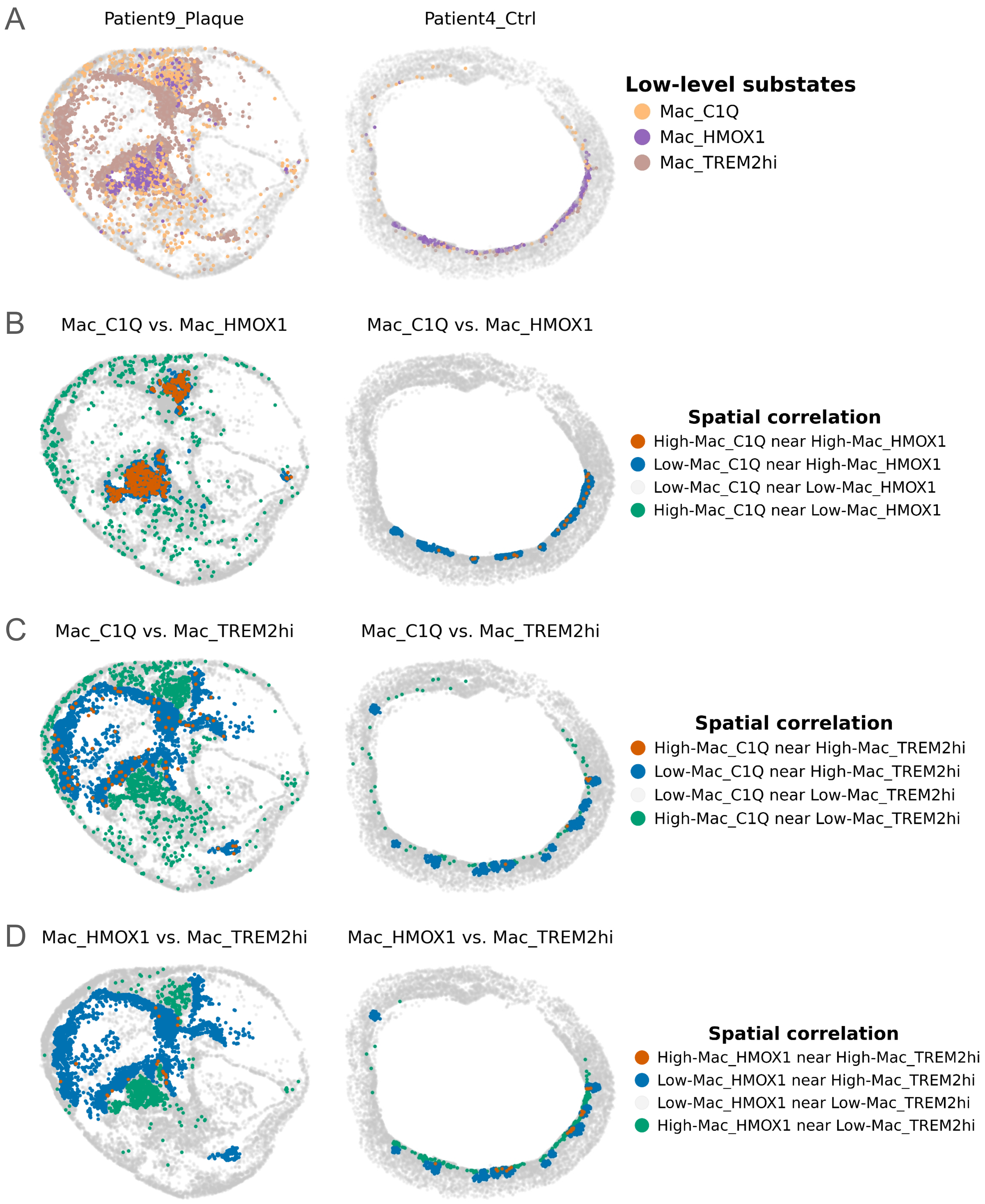


**Suppl. Fig. 27: Spatial correlation of Macrophage substates.** (A) Spatial distribution of the panel 1 Macrophage substates Mac_C1Q, Mac_HMOX1 and Mac_TREM2hi for two selected samples. (B-D) Spatial correlation calculated between substate pair combinations using Local Bivariate Moran's I statistic. Cells coloured other than grey show a significant spatial correlation across substate pairs (FDR<0.05), indicating the nature of correlation (i.e. high-high, high-low). FDR: false discovery rate, resulting from Benjamini-Hochberg correction of raw p-values of Local Bivariate Moran's I values. Multiple testing correction was performed within sample per substate pair.

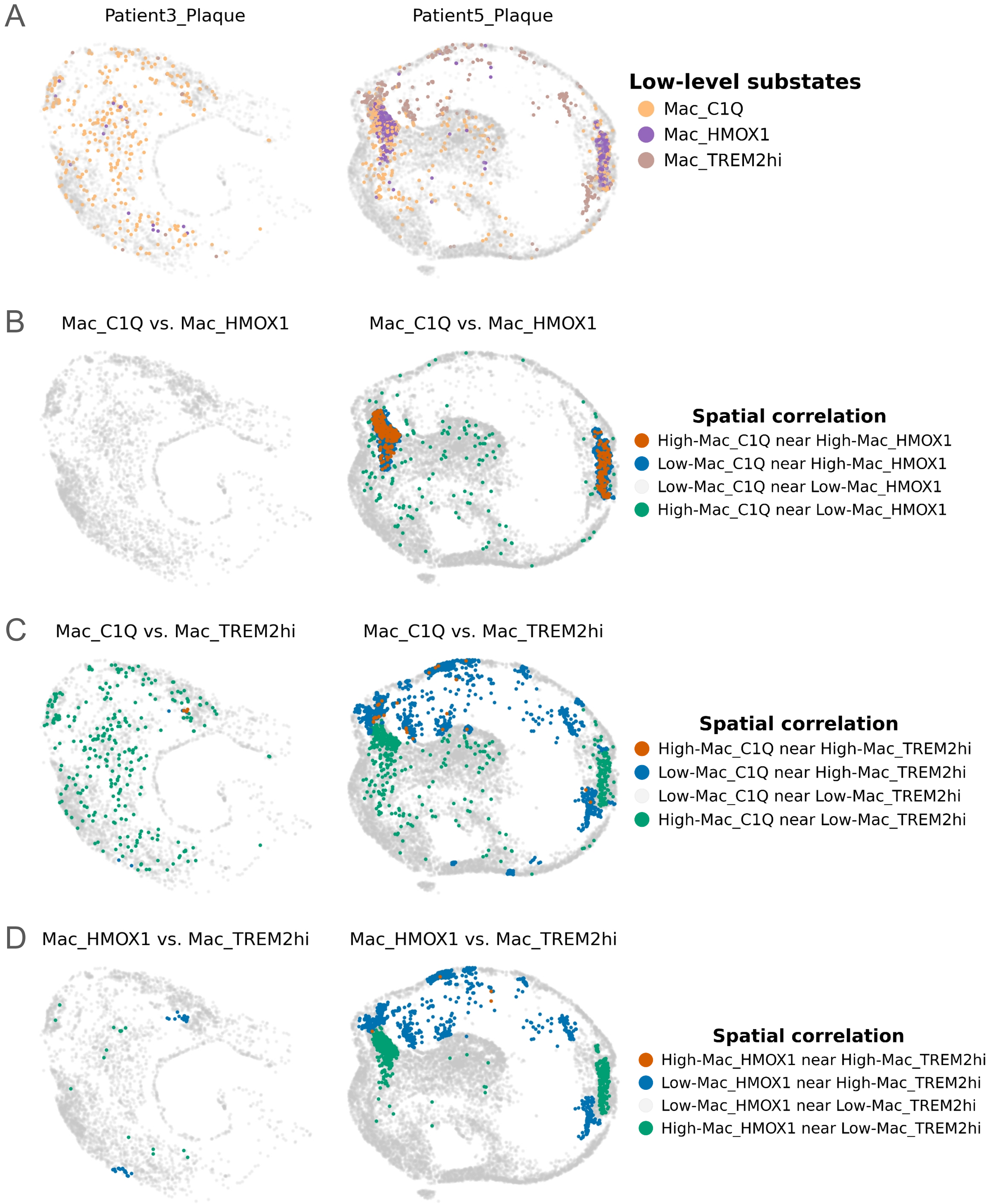


**Suppl. Fig. 28: Spatial correlation of Macrophage substates.** (A) Spatial distribution of the panel 1 Macrophage substates Mac_C1Q, Mac_HMOX1 and Mac_TREM2hi for two selected samples. (B-D) Spatial correlation calculated between substate pair combinations using Local Bivariate Moran's I statistic. Cells coloured other than grey show a significant spatial correlation across substate pairs (FDR<0.05), indicating the nature of correlation (i.e. high-high, high-low). FDR: false discovery rate, resulting from Benjamini-Hochberg correction of raw p-values of Local Bivariate Moran's I values. Multiple testing correction was performed within sample per substate pair.

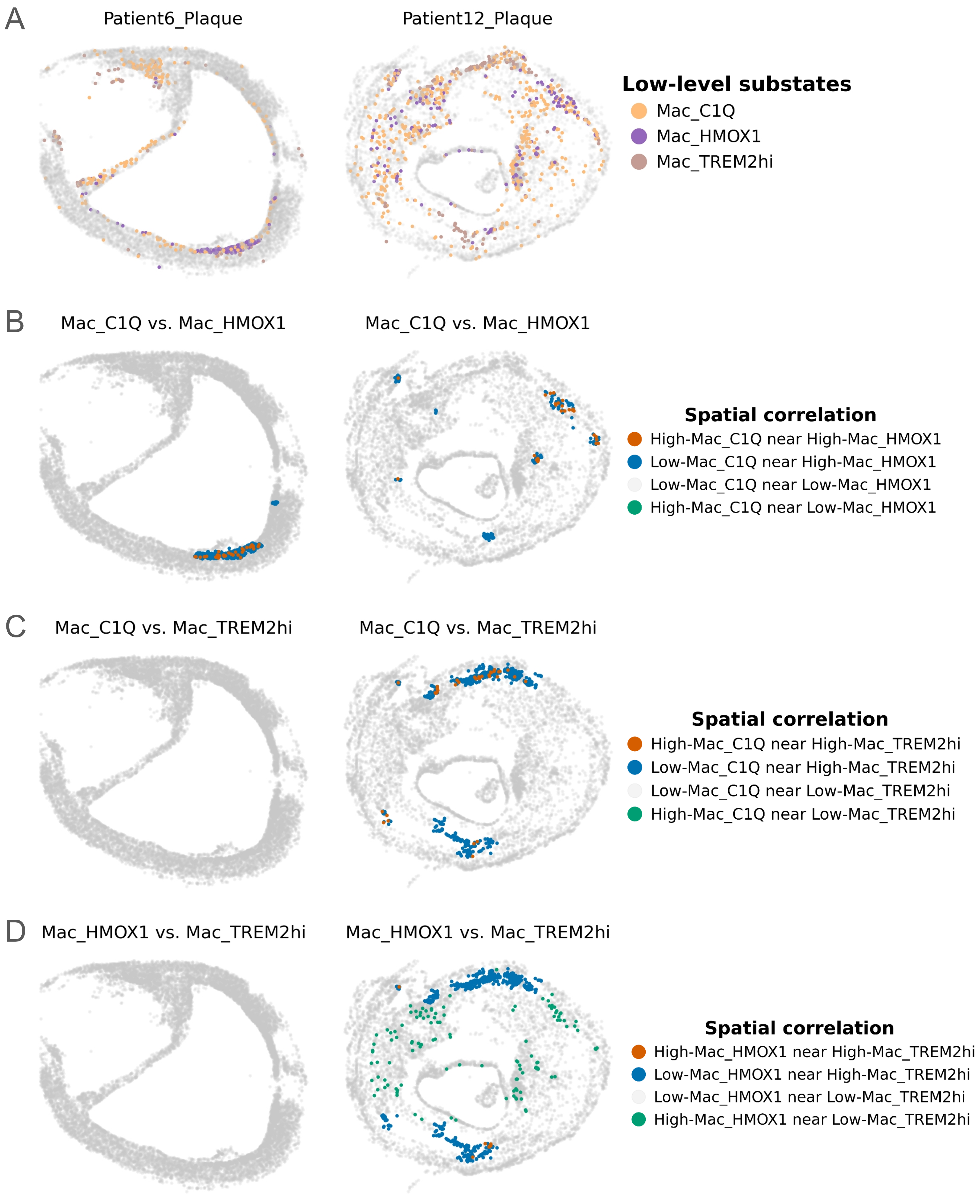


**Suppl. Fig. 29: Spatial correlation of Macrophage substates.** (A) Spatial distribution of the panel 1 Macrophage substates Mac_C1Q, Mac_HMOX1 and Mac_TREM2hi for two selected samples. (B-D) Spatial correlation calculated between substate pair combinations using Local Bivariate Moran's I statistic. Cells coloured other than grey show a significant spatial correlation across substate pairs (FDR<0.05), indicating the nature of correlation (i.e. high-high, high-low). FDR: false discovery rate, resulting from Benjamini-Hochberg correction of raw p-values of Local Bivariate Moran's I values. Multiple testing correction was performed within sample per substate pair.

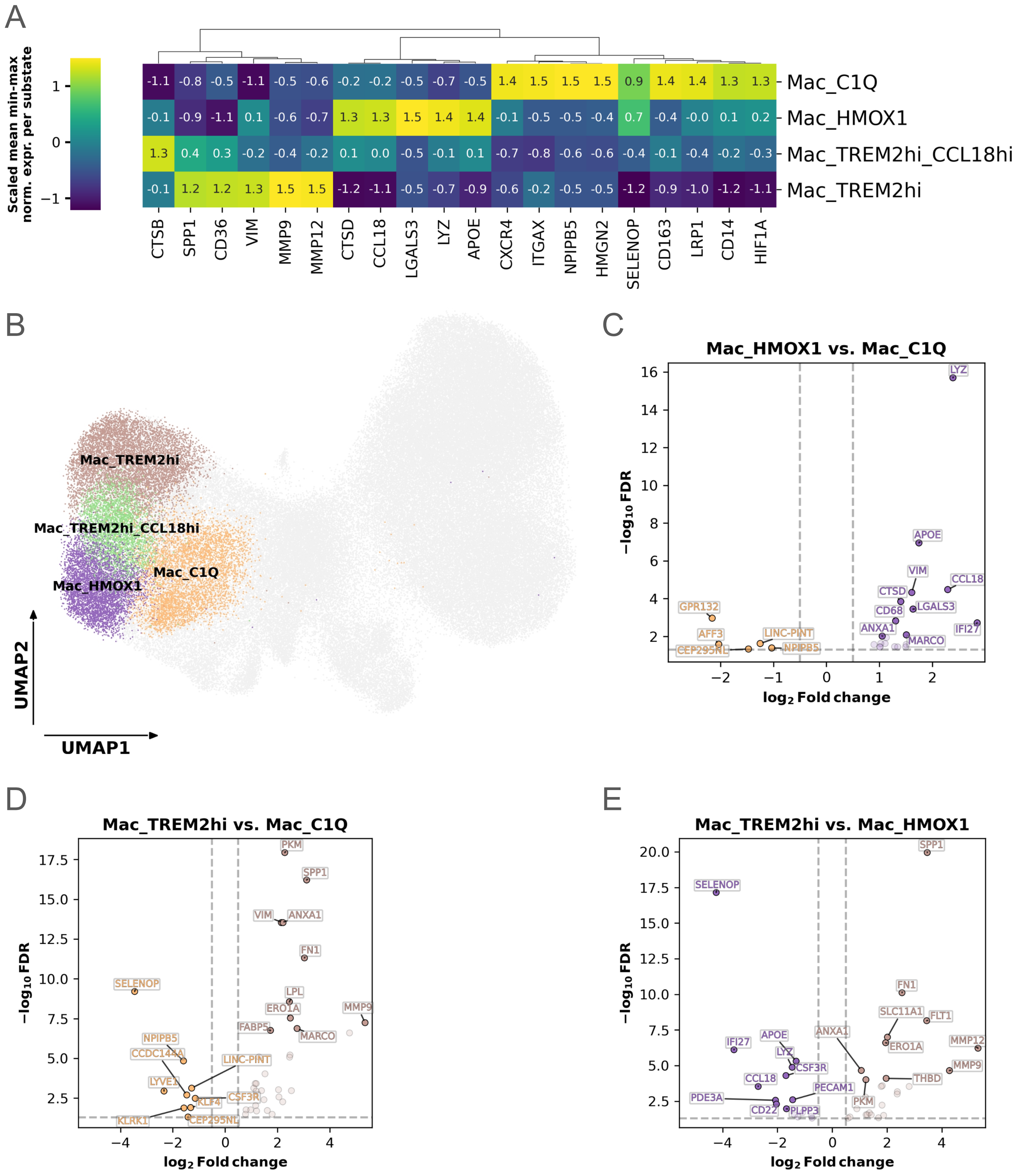


**Suppl. Fig. 30: Gene expression differences across four macrophage substates in panel 1 neighbourhood clusters 2 and 4.** Gene expression differences in Mac_C1Q, Mac_HMOX1, Mac_TREM2hi, and Mac_TREM2hi_CCL18hi cells belonging to the panel 1 neighbourhood clusters 2 and 4, coming from plaques of patients 4,7,8,10. First, gene expression was converted to the range of [0,1] by min-max normalising per sample, then averaged within cell substate. (A) Shows a clustermap of 20 genes with highest variance in the mean min-max normalised expression across substates. Clustering was done using the Z-scores of the expression mean values. (B) Low dimensional UMAP representation of cell expression with panel 1 genes, with dot colouring representing the four macrophage substates of interest. (C-E) Differential gene expression across cell substates Mac_C1Q, Mac_HMOX1, Mac_TREM2hi. X-axis depicts the log2 fold-change (log2FC) of gene expression across substates, y-axis depicts the –log10 values of the Benjamini-Hochberg adjusted significance level of gene expression change (false discovery rate, FDR). Vertical dashed lines represent the log2FC threshold at +/- 0.5, horizontal dashed line represent the FDR threshold of –log10(0.05).

A

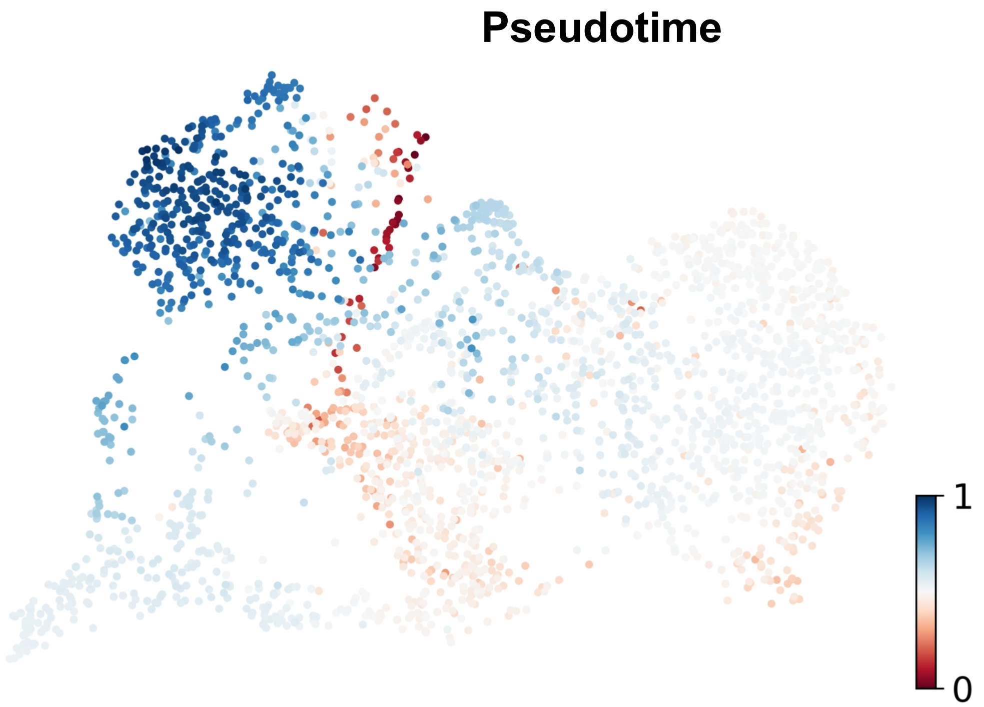


B

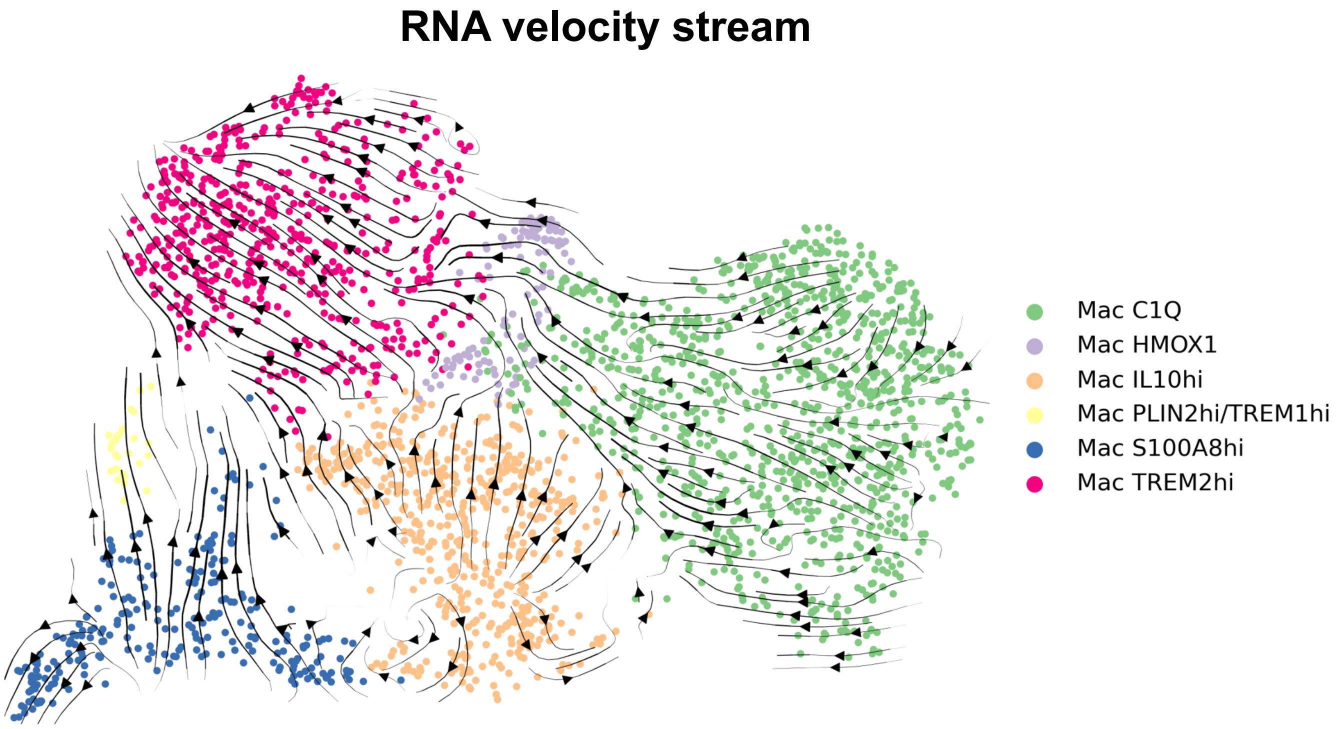


**Suppl. Fig. 31: RNA velocity of Macrophage subtypes in scRNA-seq data.** (A) Macrophage cells coloured by the normalized pseudotime inferred by TFvelo. Pseudotime reflects the progression of cell development, starting from 0 to 1. (B) RNA velocity stream plot. The arrows depict the transcriptional trajectory of the cells over pseudotime inferred by TFvelo based on cell expression data.

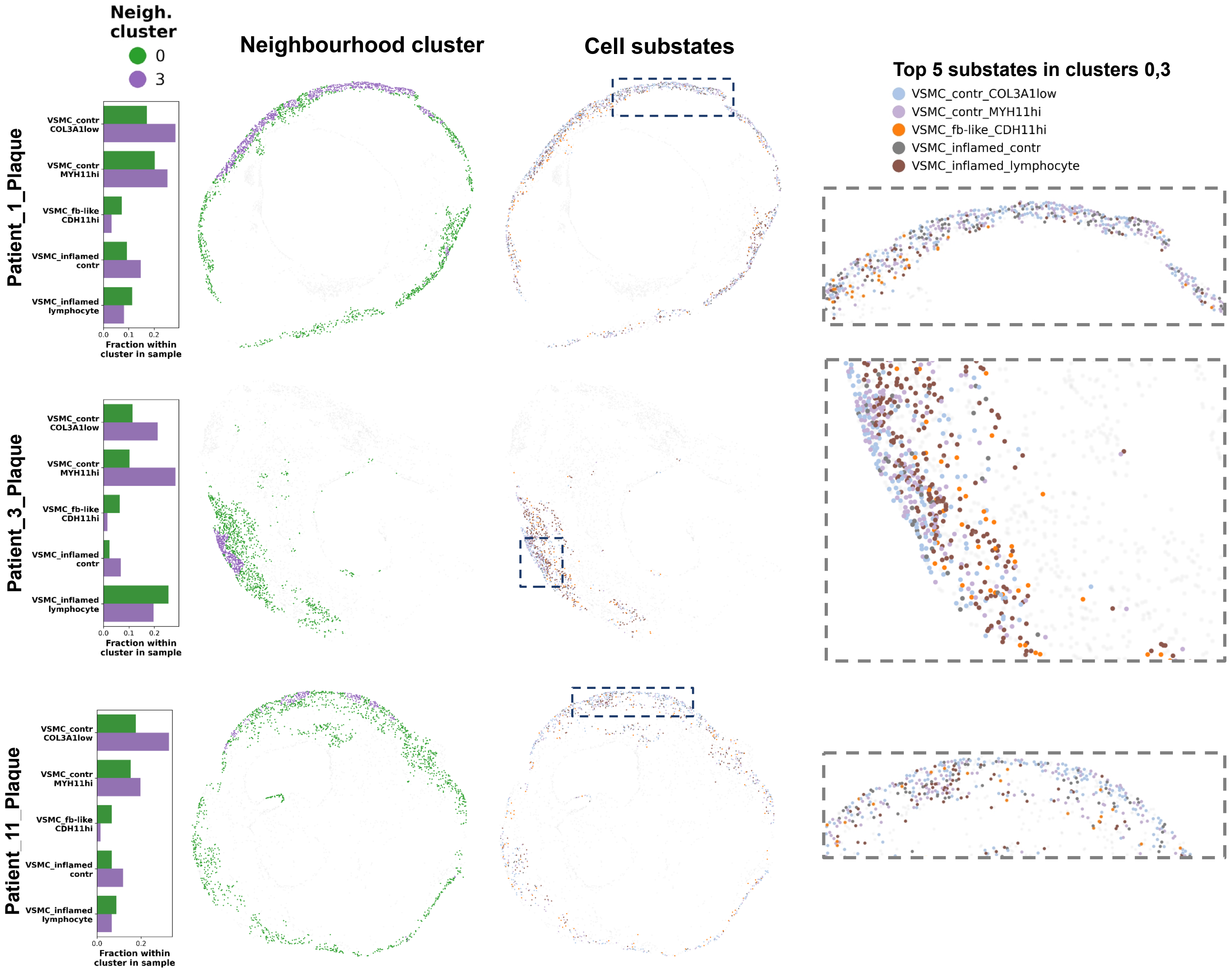


**Suppl. Fig. 32: Vascular smooth muscle cell (VSMC) neighbourhoods presented in further plaques.** Composition of VSMC neighbourhoods shown based on the five most abundant cell substates per panel 1 neighbourhood cluster, in three further samples. The first column displays relative fractions of these substates within each neighbourhood cluster per plaque. The second column shows their spatial distribution coloured by neighbourhood cluster, while the third column presents the same distribution coloured by cell substate. The fourth column provides a zoomed-in view of regions with interesting substate spatial distribution. Dashed rectangles in column three represent the original location of interesting regions within sample.

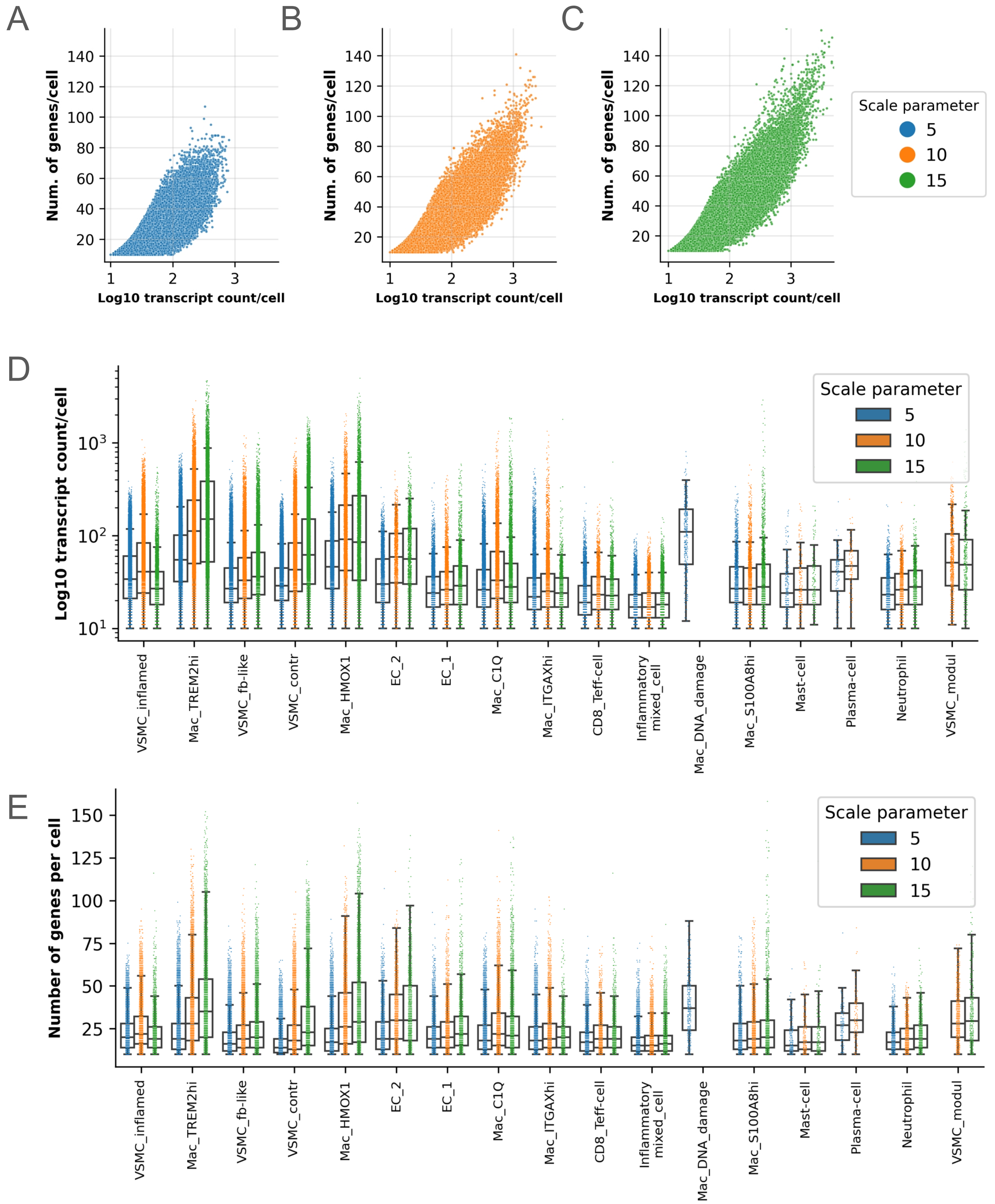


**Suppl. Fig. 33: Transcript and gene count per cell comparison of Baysor cell segmentations with different "scale" parameters on gene panel 1.** Subplots (A-C) show the number of transcript counts per cell plotted against the number of genes detected per cell, across the 3 scale parameters, with gene panel 1 data. Transcript counts per cells are plotted logarithmic. (D) Log10 transcript count per cell of low-level cell types annotated in the gene panel 1 dataset, split by "scale" parameter used for segmentation. (E) Number of genes per cell of low-level cell types annotated in the gene panel 1 dataset, split by "scale" parameter used for segmentation.

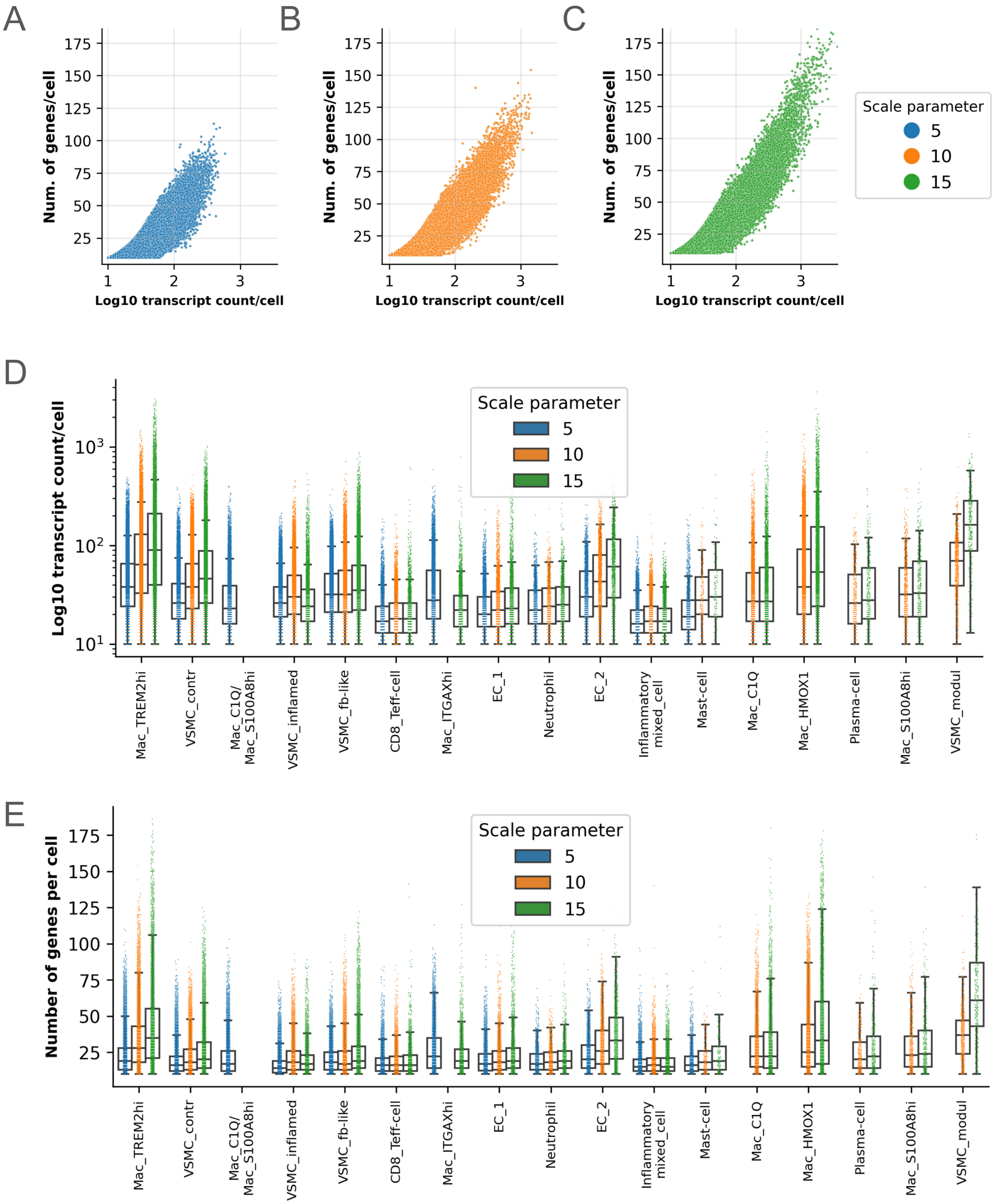


**Suppl. Fig. 34: Transcript and gene count per cell comparison of Baysor cell segmentations with different "scale" parameters on gene panel 2.** Subplots (A-C) show the number of transcript counts per cell plotted against the number of genes detected per cell, across the 3 scale parameters, with gene panel 2 data. Transcript counts per cells are plotted logarithmic. (D) Log10 transcript count per cell of low-level cell types annotated in the gene panel 2 dataset, split by "scale" parameter used for segmentation. (E) Number of genes per cell of low-level cell types annotated in the gene panel 2 dataset, split by "scale" parameter used for segmentation.

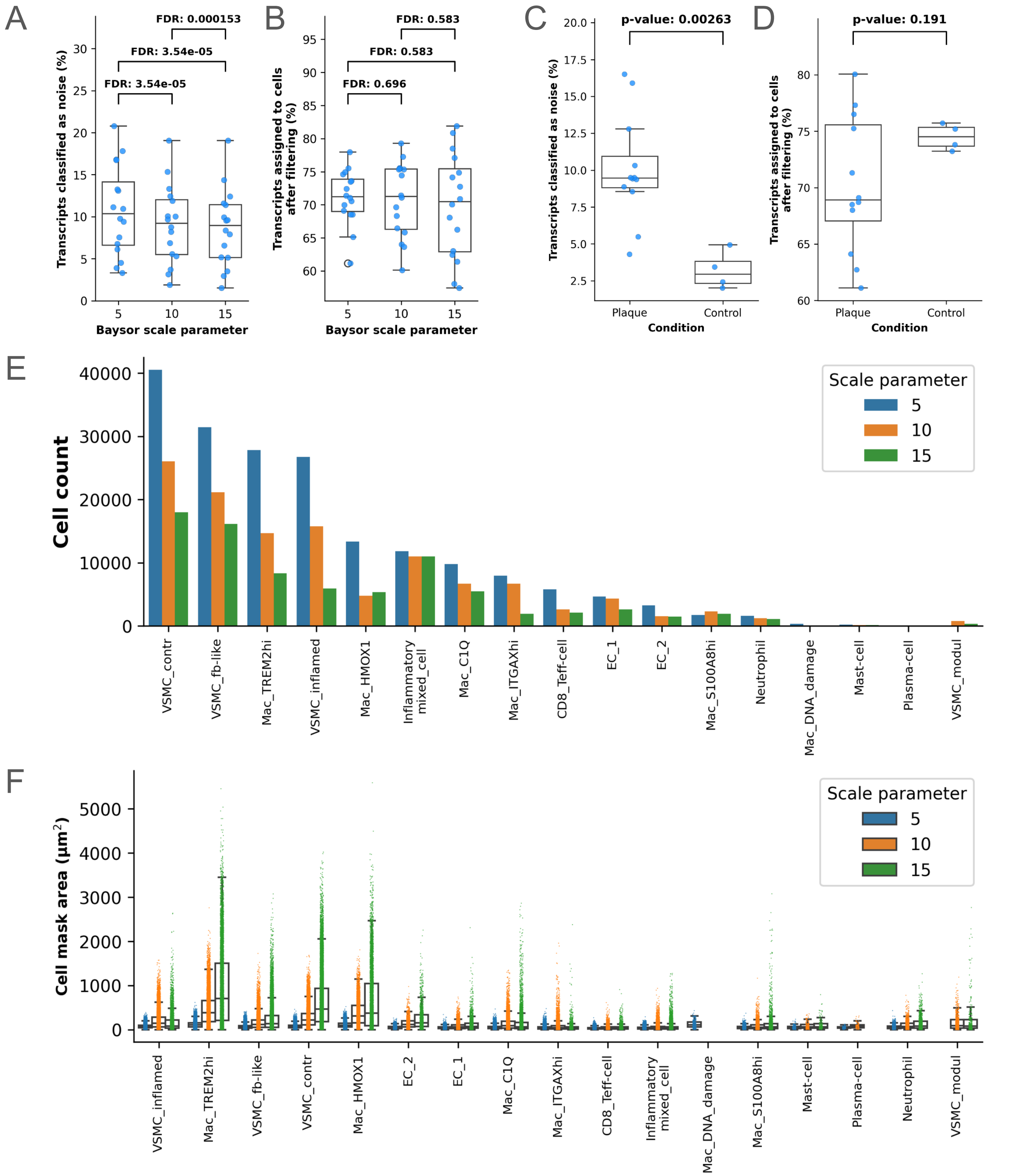


**Suppl. Fig. 35: Comparison of Baysor cell segmentations with different “scale” parameters on gene panel 1.** (A) Distribution of the percentage of transcripts classified as background noise by Baysor. Within each „scale“ parameter, dots represent the results of the best segmentation for a sample, performed with gene panel 1. Only transcripts localised inside sample area were considered. False discovery rate (FDR): Benjamini-Hochberg corrected p-values of paired t-tests performed across the “scale” parameters. (B) Distribution of the percentage of transcripts assigned to cells by Baysor. Within each „scale“ parameter, dots represent the results of the best segmentation for a sample, performed with gene panel 1. Only transcripts localised inside sample area were considered. False discovery rate (FDR): Benjamini-Hochberg corrected p-values of paired t-tests performed across the “scale” parameters. (C) Distribution of the percentage of transcripts classified as background noise by Baysor, split by condition. Dots represent the results of segmentations used for further analysis („scale“: 10 µm), performed with gene panel 1. P-value: unpaired t-test performed across the conditions. (D) Distribution of the percentage of transcripts assigned to cells by Baysor, split by condition. Dots represent the results of segmentations used for further analysis („scale“: 10 µm), performed with gene panel 1. P-value: unpaired t-test performed across the conditions. (E) Cell counts of low-level cell types annotated in the gene panel 1 dataset, split by “scale” parameter used for segmentation. (F) Cell mask areas of individual cells segmented by Baysor with gene panel 1, split by low-level cell type annotation and the “scale” parameter used for cell segmentation.

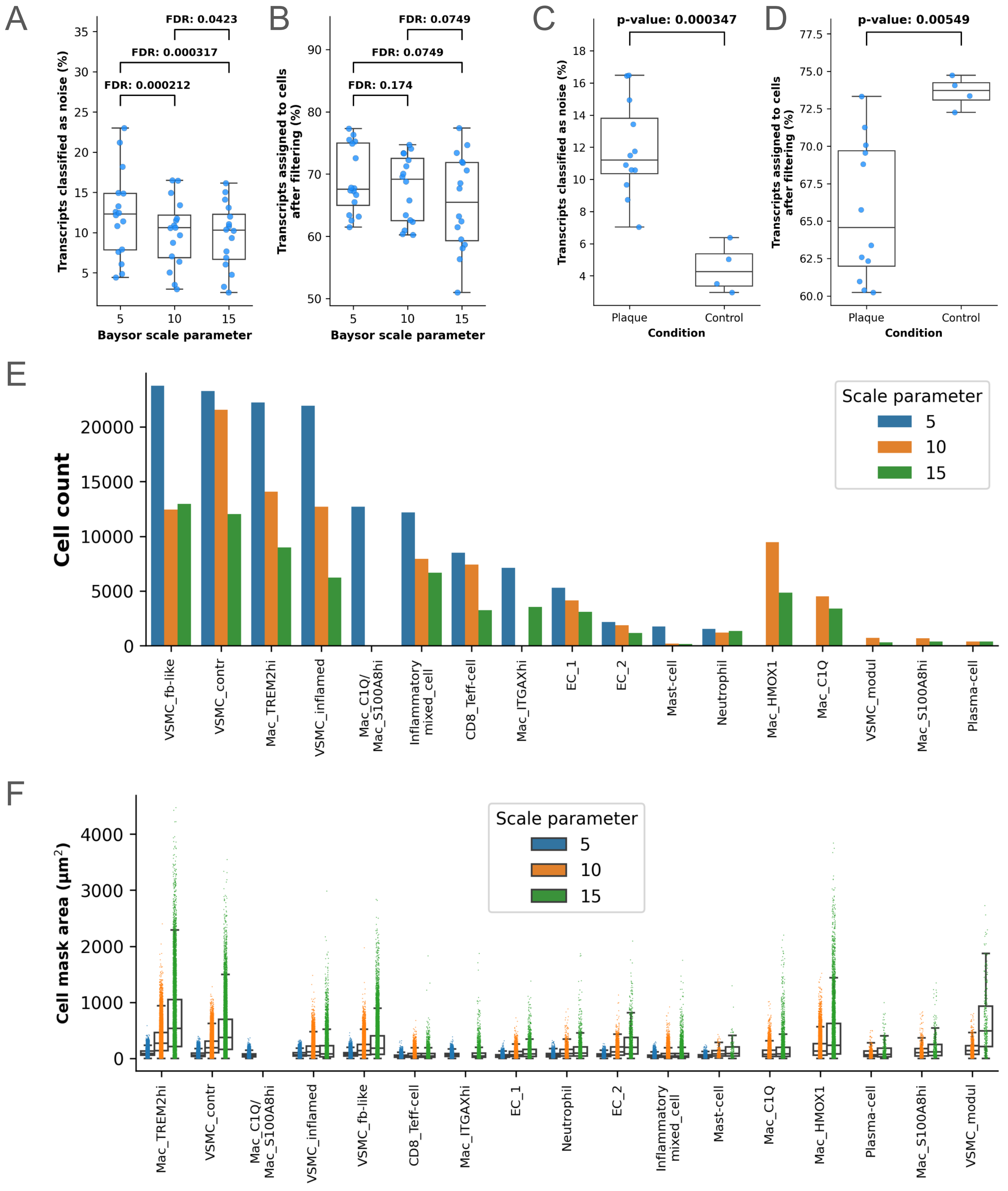


**Suppl. Fig. 36: Comparison of Baysor cell segmentations with different "scale" parameters on gene panel 2.** (A) Distribution of the percentage of transcripts classified as background noise by Baysor. Within each „scale" parameter, dots represent the results of the best segmentation for a sample, performed with gene panel 2. Only transcripts localised inside sample area were considered. False discovery rate (FDR): Benjamini-Hochberg corrected p-values of paired t-tests performed across the "scale" parameters. (B) Distribution of the percentage of transcripts assigned to cells by Baysor. Within each „scale" parameter, dots represent the results of the best segmentation for a sample, performed with gene panel 2. Only transcripts localised inside sample area were considered. False discovery rate (FDR): Benjamini-Hochberg corrected p-values of paired t-tests performed across the "scale" parameters. (C) Distribution of the percentage of transcripts classified as background noise by Baysor, split by condition. Dots represent the results of segmentations used for further analysis („scale": 10 µm), performed with gene panel 2. P-value: unpaired t-test performed across the conditions. (D) Distribution of the percentage of transcripts assigned to cells by Baysor, split by condition. Dots represent the results of segmentations used for further analysis („scale": 10 µm), performed with gene panel 2. P-value: unpaired t-test performed across the conditions. (E) Cell counts of low-level cell types annotated in the gene panel 2 dataset, split by "scale" parameter used for segmentation. (F) Cell mask areas of individual cells segmented by Baysor with gene panel 2, split by low-level cell type annotation and the "scale" parameter used for cell segmentation.

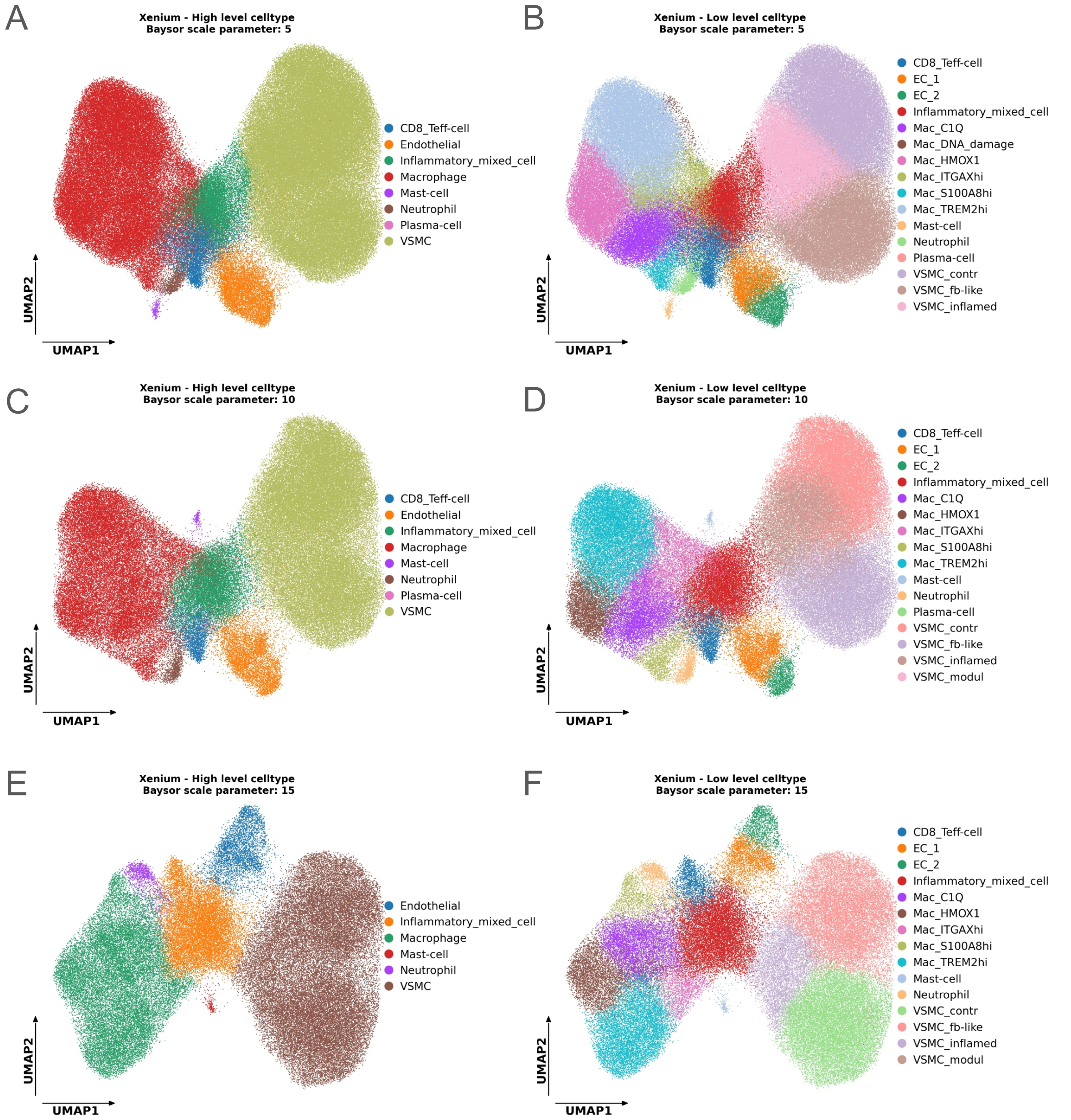


**Suppl. Fig. 37: Comparison of cell type annotations coming from cell segmentations on gene panel 1 data with different Baysor "scale" parameters.** (A) UMAP low dimensional representation of cell expression showing high-level and (B) low-level cell type annotation of cells, segmented on gene panel 1 data with Baysor "scale" parameter 5. (C) UMAP low dimensional representation of cell expression showing high-level and (D) low-level cell type annotation of cells, segmented on gene panel 1 data with Baysor "scale" parameter 10. (E) UMAP low dimensional representation of cell expression showing high-level and (F) low-level cell type annotation of cells, segmented on gene panel 1 data with Baysor "scale" parameter 5. UMAP: Uniform manifold maximum approximation and projection

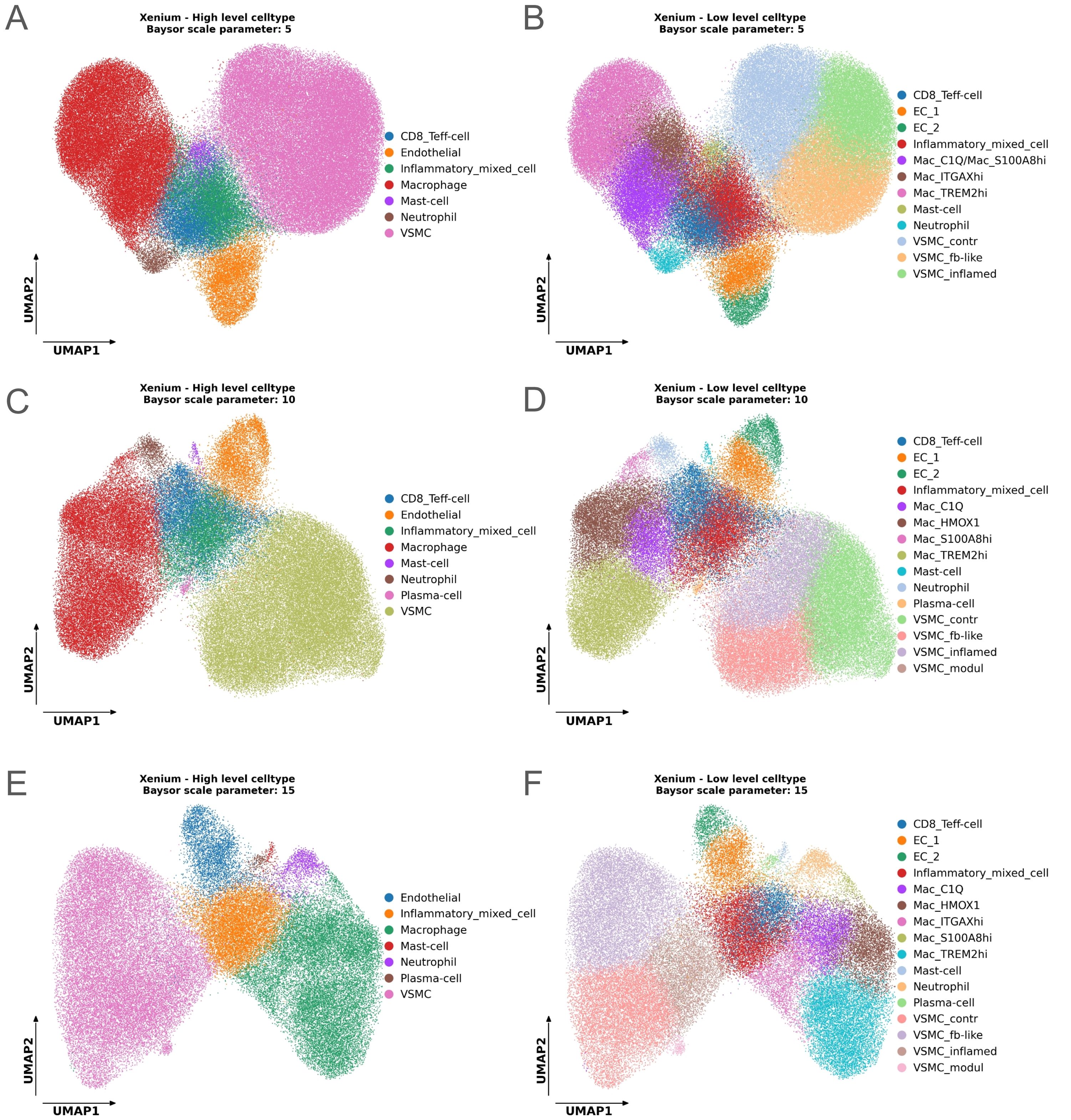


**Suppl. Fig. 38: Comparison of cell type annotations coming from cell segmentations on gene panel 2 data with different Baysor "scale" parameters.** (A) UMAP low dimensional representation of cell expression showing high-level and (B) low-level cell type annotation of cells, segmented on gene panel 2 data with Baysor "scale" parameter 5. (C) UMAP low dimensional representation of cell expression showing high-level and (D) low-level cell type annotation of cells, segmented on gene panel 2 data with Baysor "scale" parameter 10. (E) UMAP low dimensional representation of cell expression showing high-level and (F) low-level cell type annotation of cells, segmented on gene panel 2 data with Baysor "scale" parameter 5. UMAP: Uniform manifold maximum approximation and projection

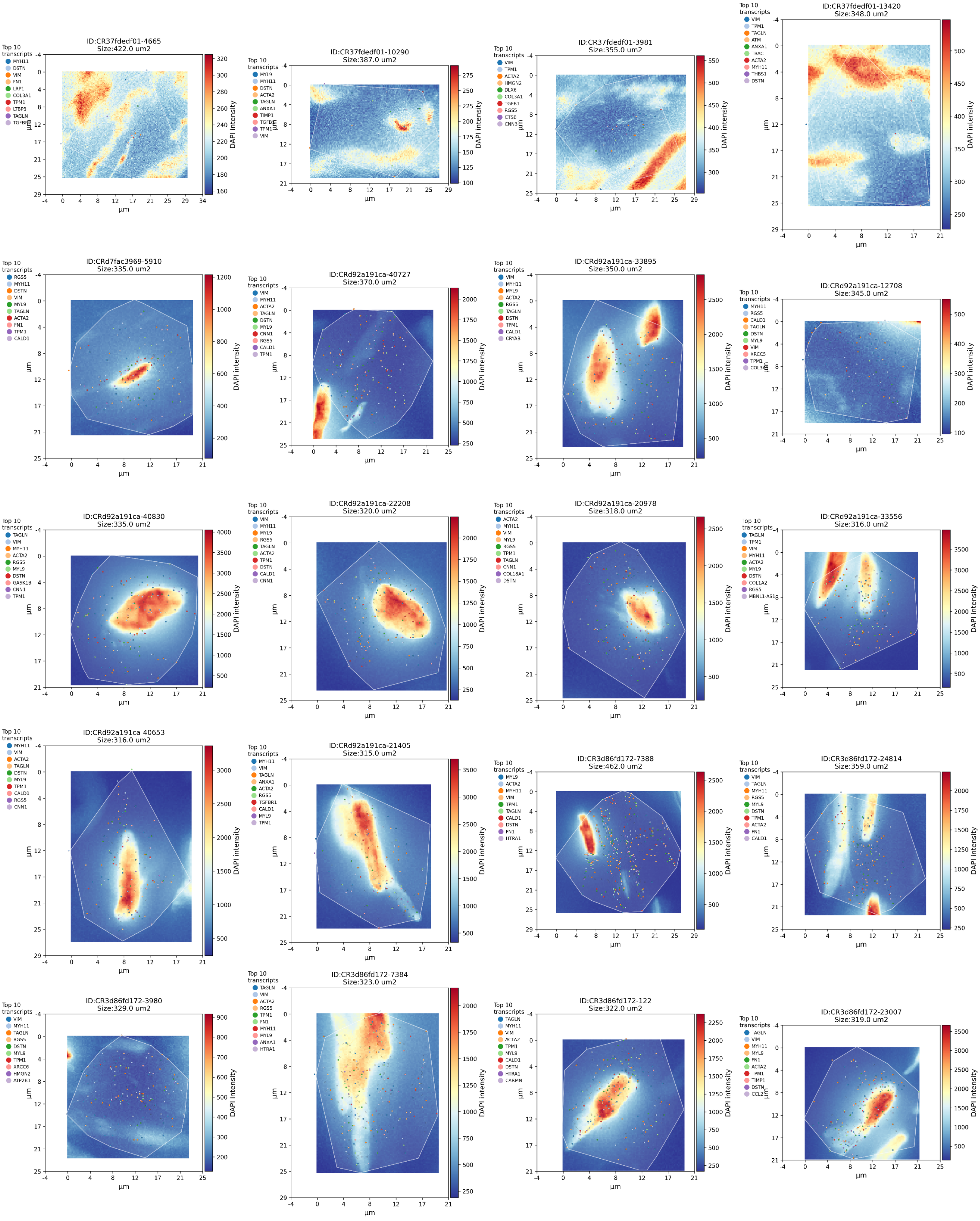

**Suppl. Fig. 39: 20 largest contractile vascular smooth muscle cells (VSMC_contr) segmented with Baysor scale parameter of 5 µm, gene panel 1.** The subplots show the 20 largest VSMC_contr cells segmented with Baysor scale parameter of 5 µm with gene panel 1 genes. The pixel intensities reflect the DAPI-staining intensity, the individual coloured dots show the 10 most abundant RNA transcripts found in the cell and the white shaded polygon displays the cell area as segmented by Baysor. The subplot title contains the individual cell-id and the cell size.

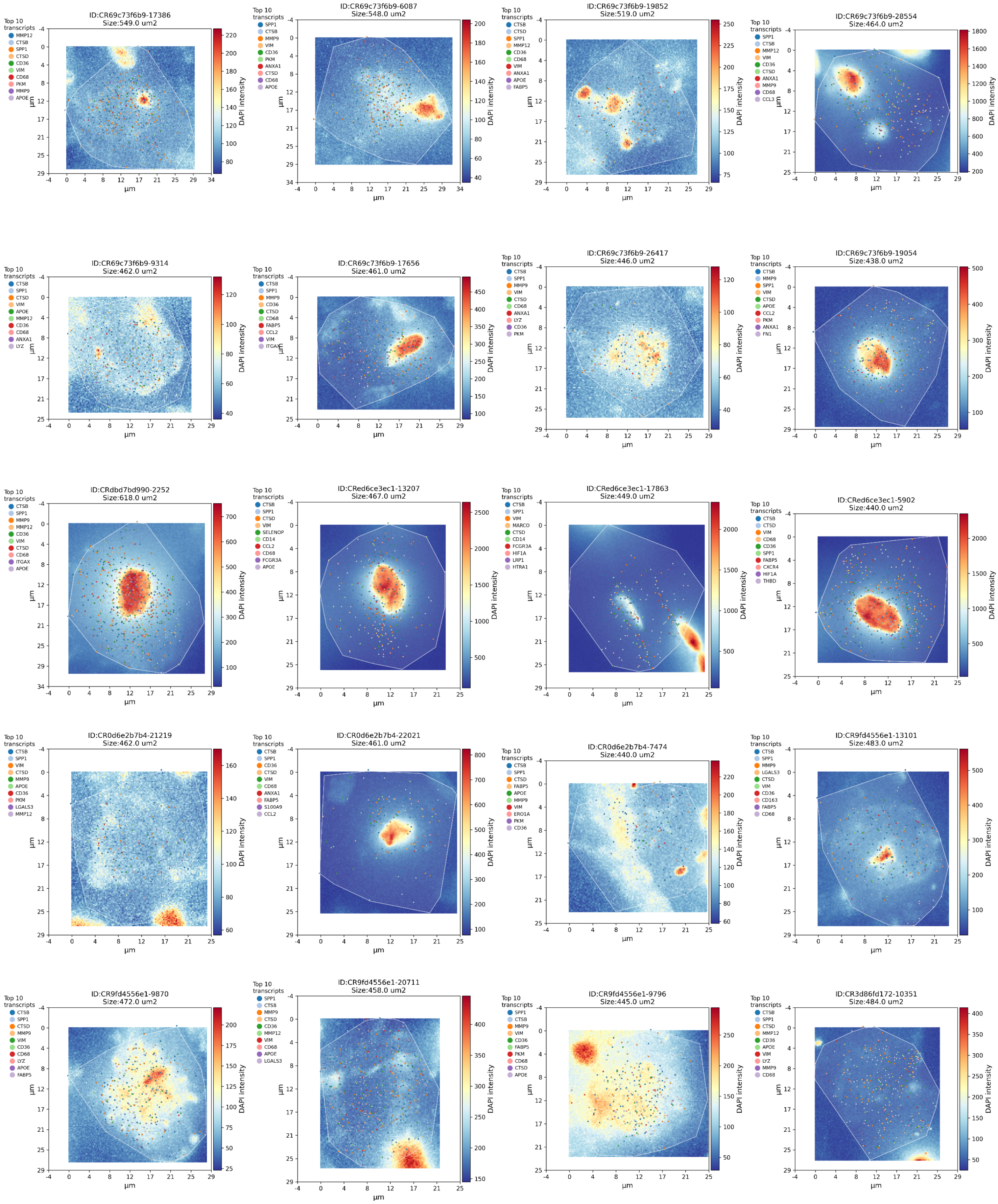


**Suppl. Fig. 40: 20 largest macrophages with high TREM2 expression (Mac_TREM2hi) segmented with Baysor scale parameter of 5 µm, gene panel 1.** The subplots show the 20 largest Mac_TREM2hi cells segmented with Baysor scale parameter of 5 µm with gene panel 1 genes. The pixel intensities reflect the DAPI-staining intensity, the individual coloured dots show the 10 most abundant RNA transcripts found in the cell and the white shaded polygon displays the cell area as segmented by Baysor. The subplot title contains the individual cell-id and the cell size.

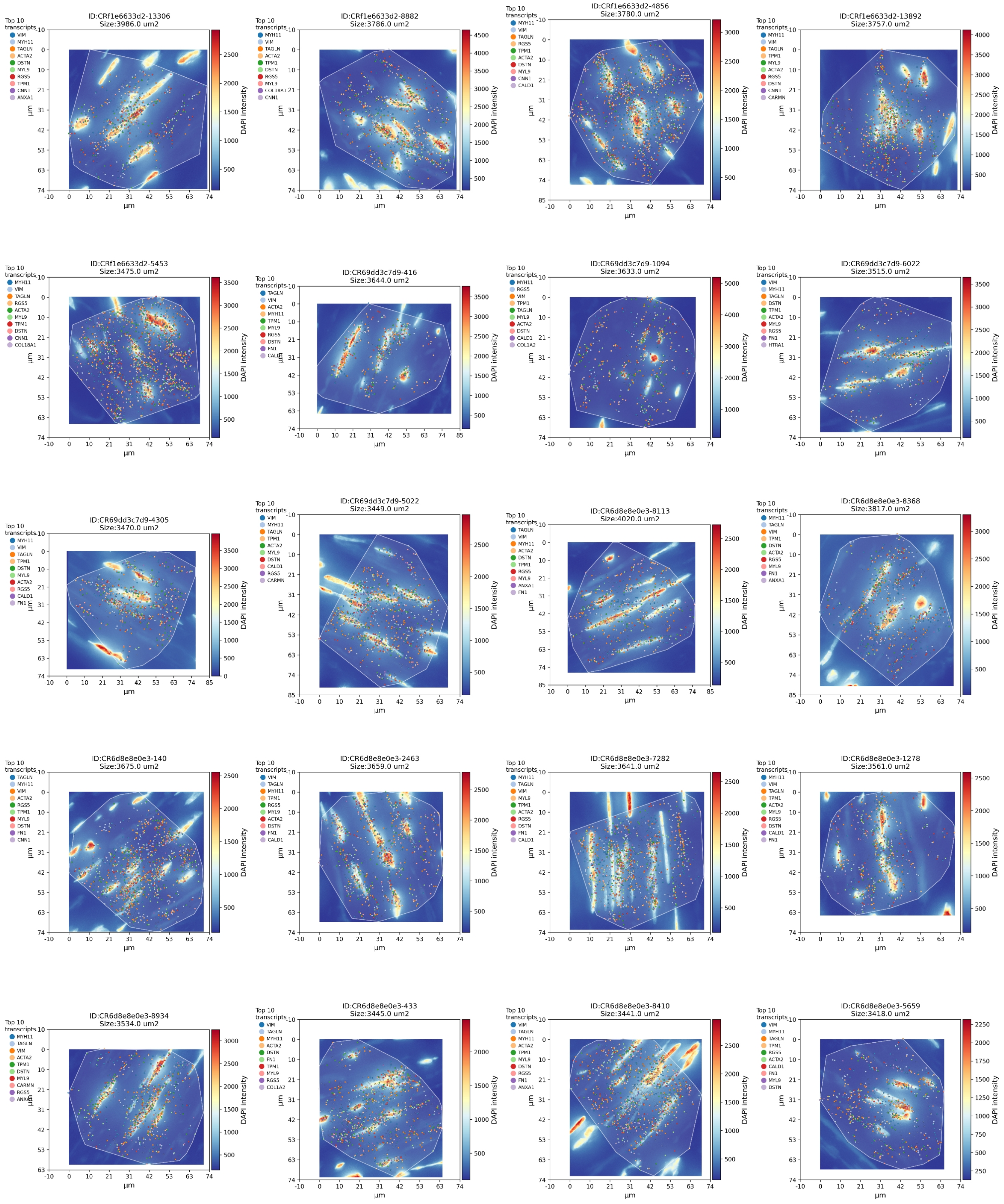


**Suppl. Fig. 41: 20 largest contractile vascular smooth muscle cells (VSMC_contr) segmented with Baysor scale parameter of 15 µm, gene panel 1.** The subplots show the 20 largest VSMC_contr cells segmented with Baysor scale parameter of 15 µm with gene panel 1 genes. The pixel intensities reflect the DAPI-staining intensity, the individual coloured dots show the 10 most abundant RNA transcripts found in the cell and the white shaded polygon displays the cell area as segmented by Baysor. The subplot title contains the individual cell-id and the cell size.

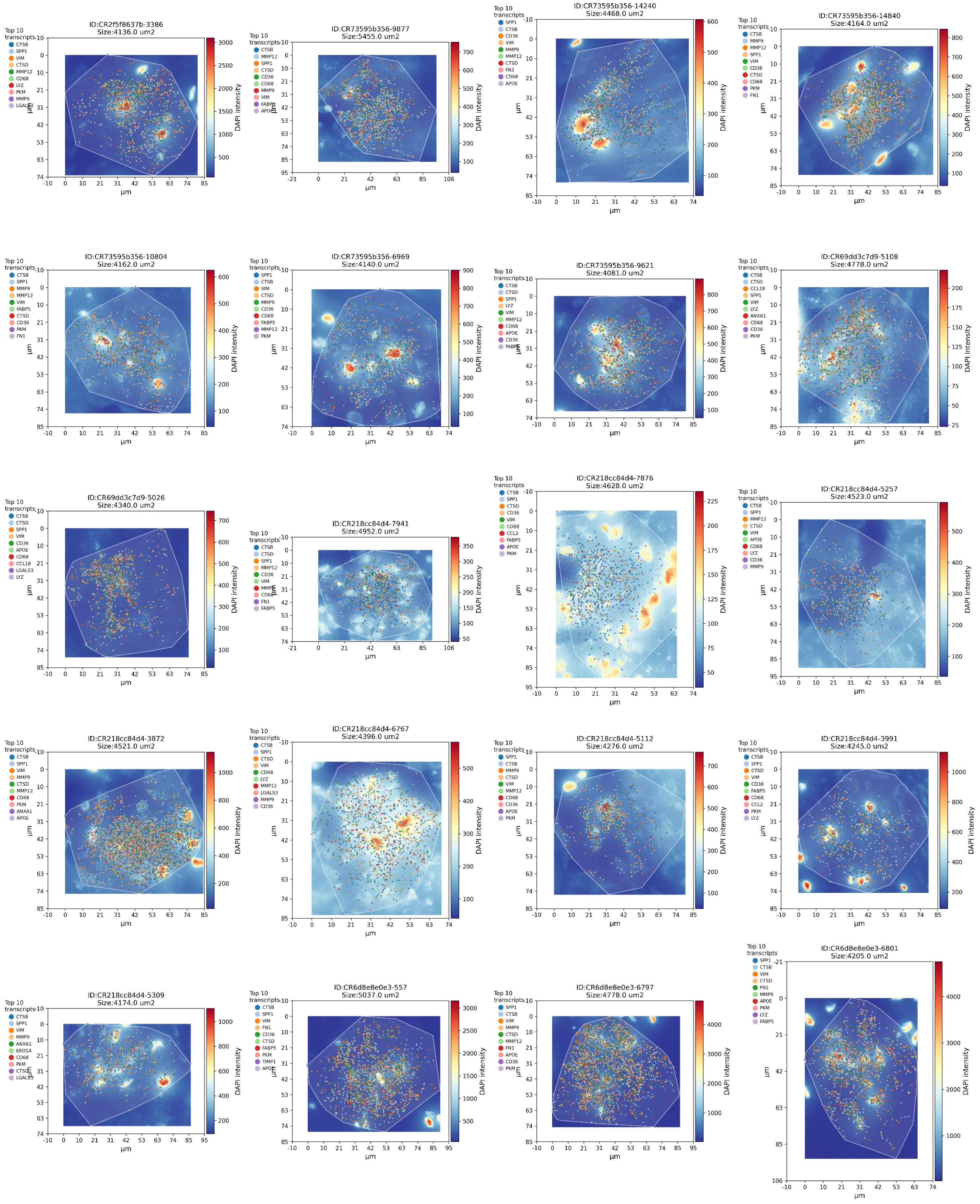


**Suppl. Fig. 42: 20 largest macrophages with high TREM2 expression (Mac_TREM2hi) segmented with Baysor scale parameter of 15 µm, gene panel 1.** The subplots show the 20 largest Mac_TREM2hi cells segmented with Baysor scale parameter of 15 µm with gene panel 1 genes. The pixel intensities reflect the DAPI-staining intensity, the individual coloured dots show the 10 most abundant RNA transcripts found in the cell and the white shaded polygon displays the cell area as segmented by Baysor. The subplot title contains the individual cell-id and the cell size.

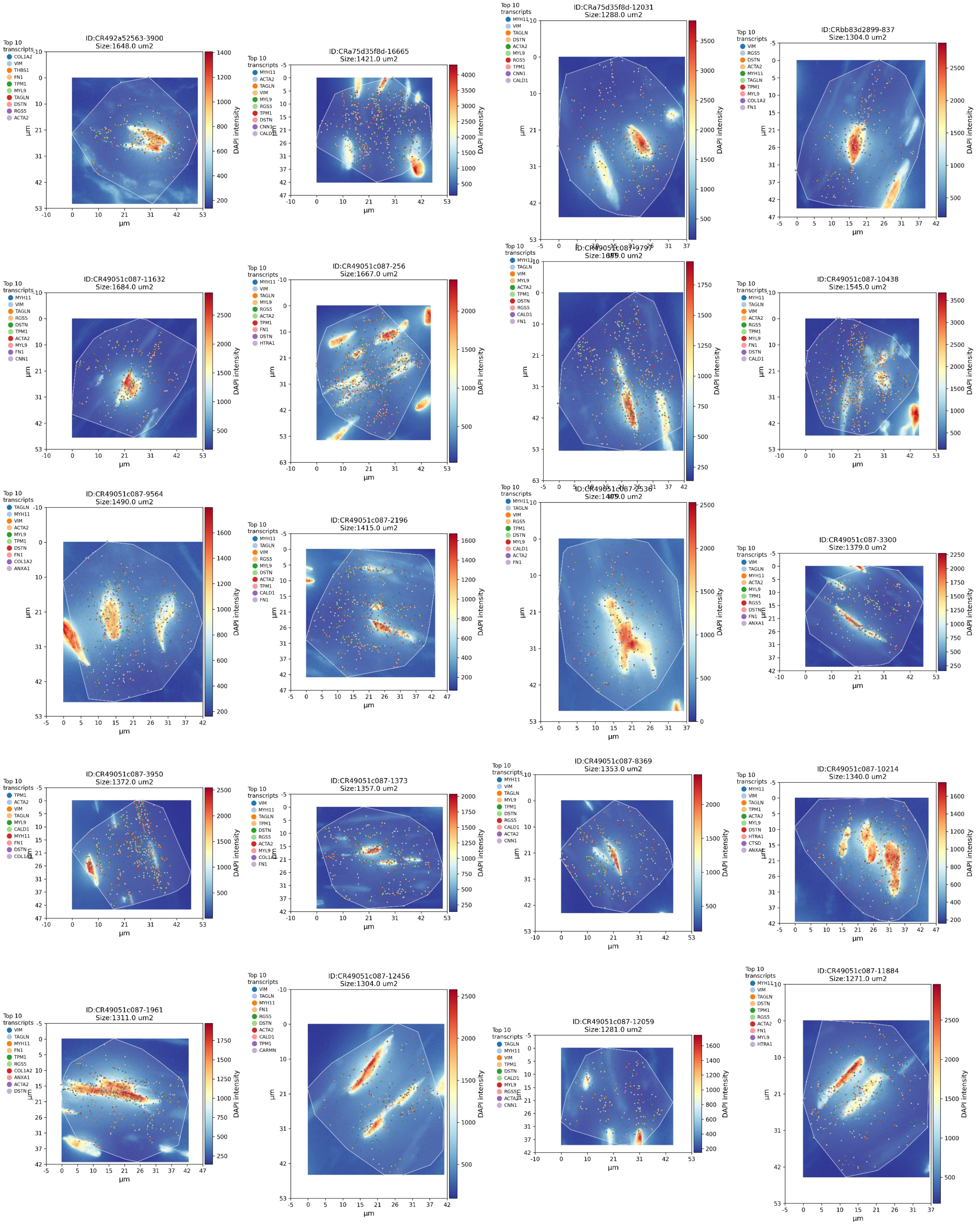


**Suppl. Fig. 43: 20 largest contractile vascular smooth muscle cells (VSMC_contr) segmented with Baysor scale parameter of 10 µm, gene panel 1.** The subplots show the 20 largest VSMC_contr cells segmented with Baysor scale parameter of 10 µm with gene panel 1 genes. The pixel intensities reflect the DAPI-staining intensity, the individual coloured dots show the 10 most abundant RNA transcripts found in the cell and the white shaded polygon displays the cell area as segmented by Baysor. The subplot title contains the individual cell-id and the cell size.

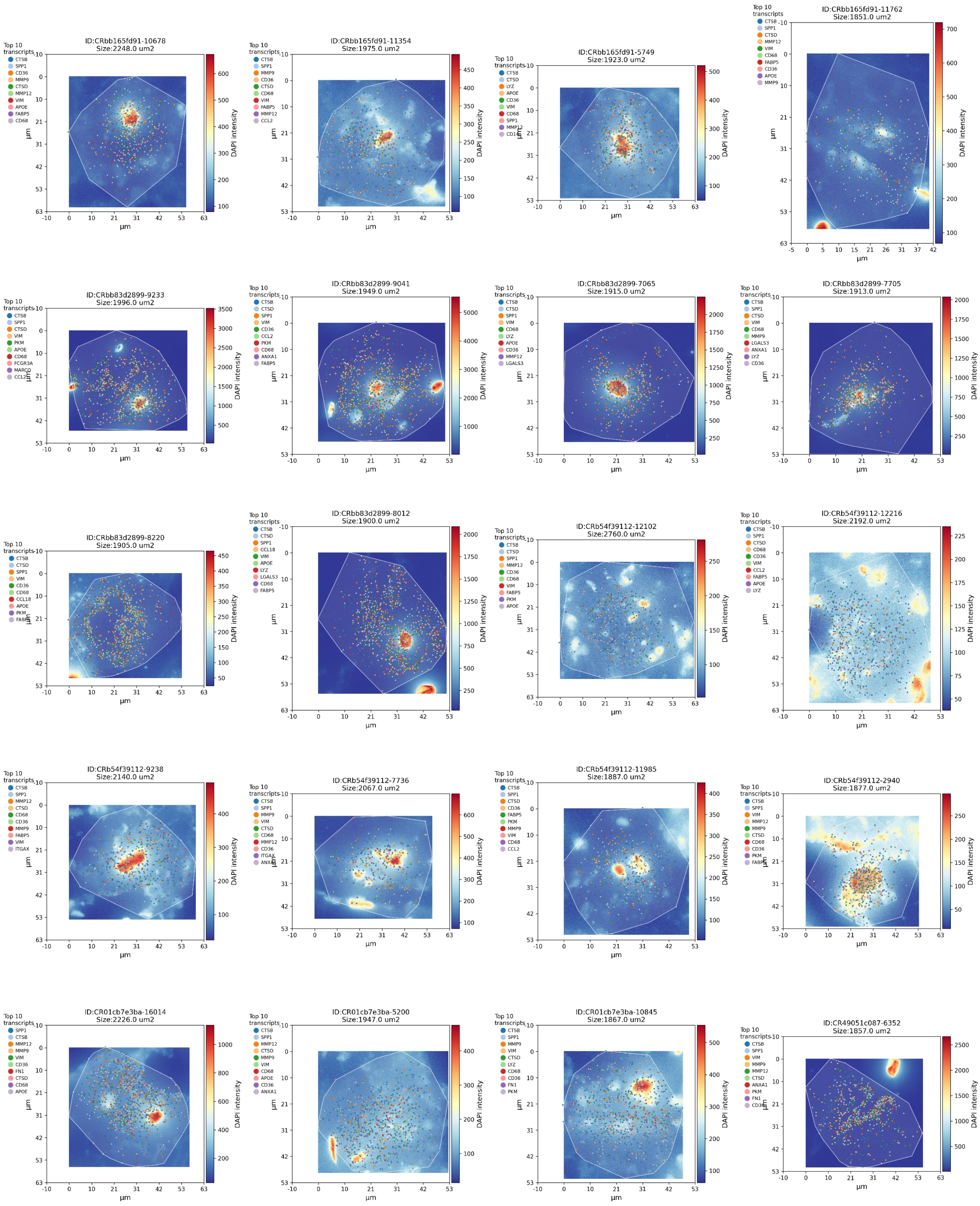


**Suppl. Fig. 44: 20 largest macrophages with high TREM2 expression (Mac_TREM2hi) segmented with Baysor scale parameter of 10 µm, gene panel 1.** The subplots show the 20 largest Mac_TREM2hi cells segmented with Baysor scale parameter of 10 µm with gene panel 1 genes. The pixel intensities reflect the DAPI-staining intensity, the individual coloured dots show the 10 most abundant RNA transcripts found in the cell and the white shaded polygon displays the cell area as segmented by Baysor. The subplot title contains the individual cell-id and the cell size.

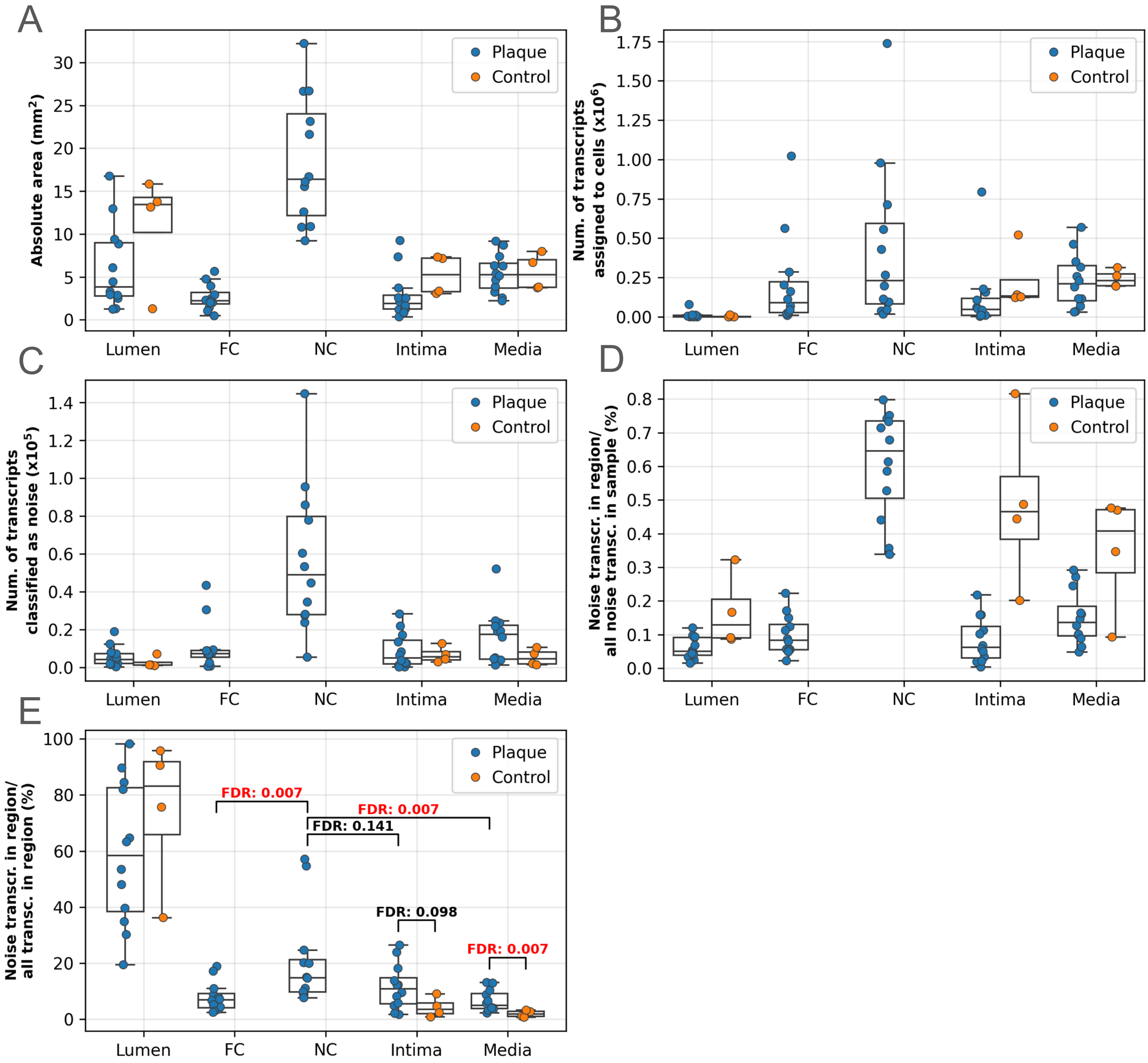


**Suppl. Fig. 45: Distribution of noise in the panel 1 segmentations used for final analysis, split by sample subregions.** The subplots show the distribution of the absolute area and signal-to-noise metrics of the panel 1 segmentations used for analysis (scale parameter: 10 µm), split by sample subregions. Each dot represents a sample, dot colouring refers to sample condition (plaque or control). (A) Absolute subregion areas of samples. (B) Absolute number of transcripts assigned to cells. (C) Absolute number of transcripts classified as noise. (D) Number of transcripts classified as noise within each subregion normalised to all noise transcripts in sample. (E) Number of transcripts classified as noise within each subregion normalised to all transcripts in within the given subregion. Unpaired Wilcoxon-test were performed to compare plaque-control distributions in the case of lumen, intima and media subregions. FDR: false discovery rate, Benjamini-Hochberg corrected p-values. Only transcripts localised on sample area were considered. FC: fibrous cap, NC: necrotic core.

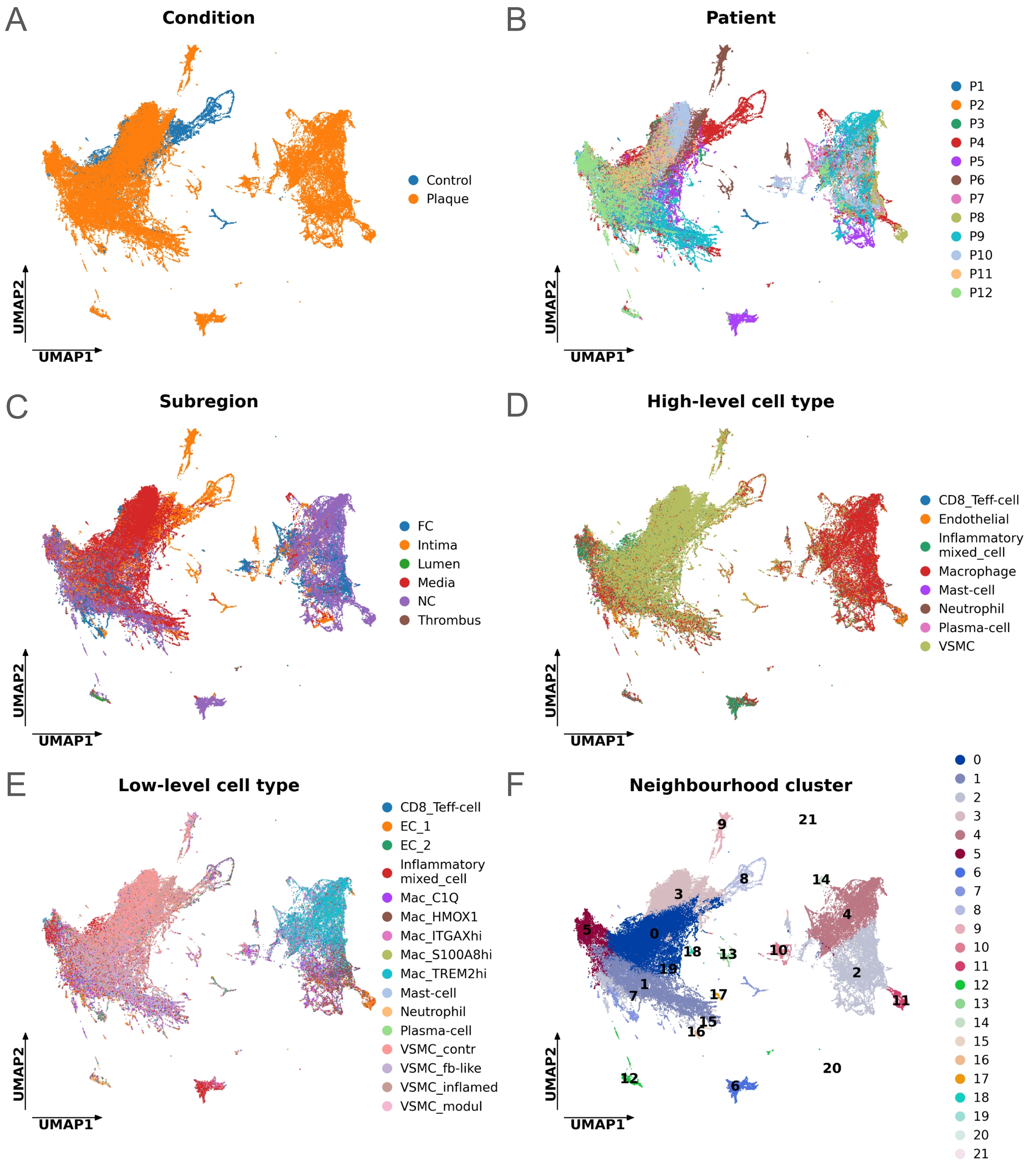


**Suppl. Fig. 46: Low dimensional UMAP representation of gene panel 1 neighbourhood matrix coloured by different cell metadata.** UMAP low dimensional representation of cell neighbourhood matrix constructed on gene panel 1 data, coloured by (A) condition, (B) patient, (C) subregion, (D) high-level cell type, (E) low-level cell type, (F) neighbourhood clusters, resulting from Leiden-clustering performed on the neighbourhood matrix. UMAP: Uniform manifold maximum approximation and projection.

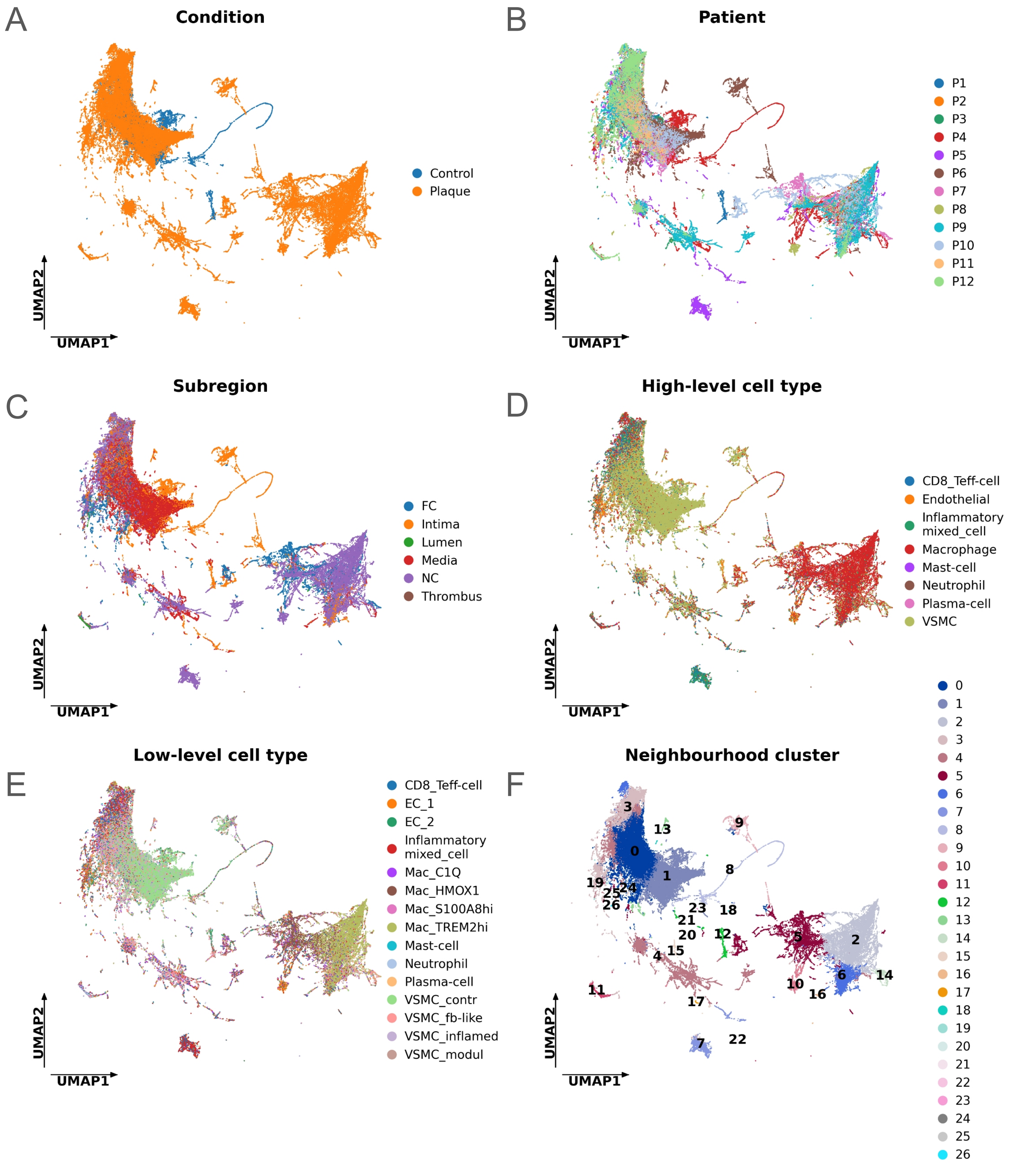


**Suppl. Fig. 47: Low dimensional UMAP representation of gene panel 2 neighbourhood matrix coloured by different cell metadata.** UMAP low dimensional representation of cell neighbourhood matrix constructed on gene panel 2 data, coloured by (A) condition, (B) patient, (C) subregion, (D) high-level cell type, (E) low-level cell type, (F) neighbourhood clusters, resulting from Leiden-clustering performed on the neighbourhood matrix. UMAP: Uniform manifold maximum approximation and projection.
